# An LSEC-focused computational drug repurposing platform for liver fibrosis: Identification of vorinostat and other LSEC-protective candidates

**DOI:** 10.64898/2026.05.23.727430

**Authors:** Rui Zuo, Mi Wang, Yun-Ting Wang, Jenny Z. Hu, Alexandra K. Moura, Dan Wang, Pin-Lan Li, Mingfu Wu, Tahir Hussain, Wei Gao, Xiang Li, Yang Zhang

**Affiliations:** Department of Pharmacological and Pharmaceutical Sciences, College of Pharmacy, University of Houston, Houston, TX, USA; Heart and Kidney Institute, College of Pharmacy, University of Houston, Houston, TX, USA; Department of Gastroenterology and Hepatology, Tongji Hospital, Tongji Medical College, Huazhong University of Science and Technology, Wuhan, Hubei, China; Department of Pharmacology and Toxicology, Virginia Commonwealth University, School of Medicine, Richmond, VA, USA

**Author notes:** **Correspondence to**: Yang Zhang, PhD, Department of Pharmacological & Pharmaceutical Sciences, Heart and Kidney Institute, College of Pharmacy, University of Houston, Houston, TX 77204-5056. These authors contributed equally to the work (Rui Zuo, Mi Wang).

**Keywords:** Liver Fibrosis, Liver Sinusoidal Endothelial Cells (LSECs), Computational Drug Screening, Drug Repurposing, Vorinostat, Hepatocyte-to-LSEC crosstalk

## Abstract

Liver sinusoidal endothelial cells (LSECs) are increasingly recognized as a critical yet underexplored cell type in anti-fibrotic drug development. This study presents a computational drug screening platform integrating LSEC-specific transcriptomic analysis across simple steatosis, fibrotic nonalcoholic steatohepatitis (NASH), and cirrhosis, with tiered gene signature selection combining machine learning, large language model-assisted curation, gene safety assessment, and Connectivity Map-based screening using human endothelial perturbational profiles. The platform identifies 6 clinical-stage and 8 preclinical candidates with LSEC-protective potential. Among these, vorinostat (SAHA), a clinically approved histone deacetylase (HDAC) inhibitor, is selected for experimental validation. In hepatocyte-specific Asah1-deficient mice fed a Paigen diet, SAHA attenuates hepatic inflammation, fibrosis, LSEC dysfunction, and portal hemodynamic abnormalities, with effects confirmed in a hepatotoxin (CCl_4_)-induced fibrosis model. High mobility group box 1 (HMGB1) is identified as a key hepatocyte-derived paracrine mediator of LSEC injury through Transwell co-culture and glycyrrhizin rescue. Vorinostat dose-dependently reverses HMGB1-induced LSEC dysfunction across inflammation, capillarization, fibrogenesis, and vasoconstriction, associated with endothelial transcription factor reprogramming including KLF2 upregulation, validated in primary LSECs and in vivo. SAHA also protected LSECs from TNF-α-induced inflammation and reduced monocyte adhesion. These findings establish an LSEC-focused drug repurposing framework and identify candidates for LSEC-protective anti-fibrotic therapy.

**GRAPHICAL ABSTRACT:** 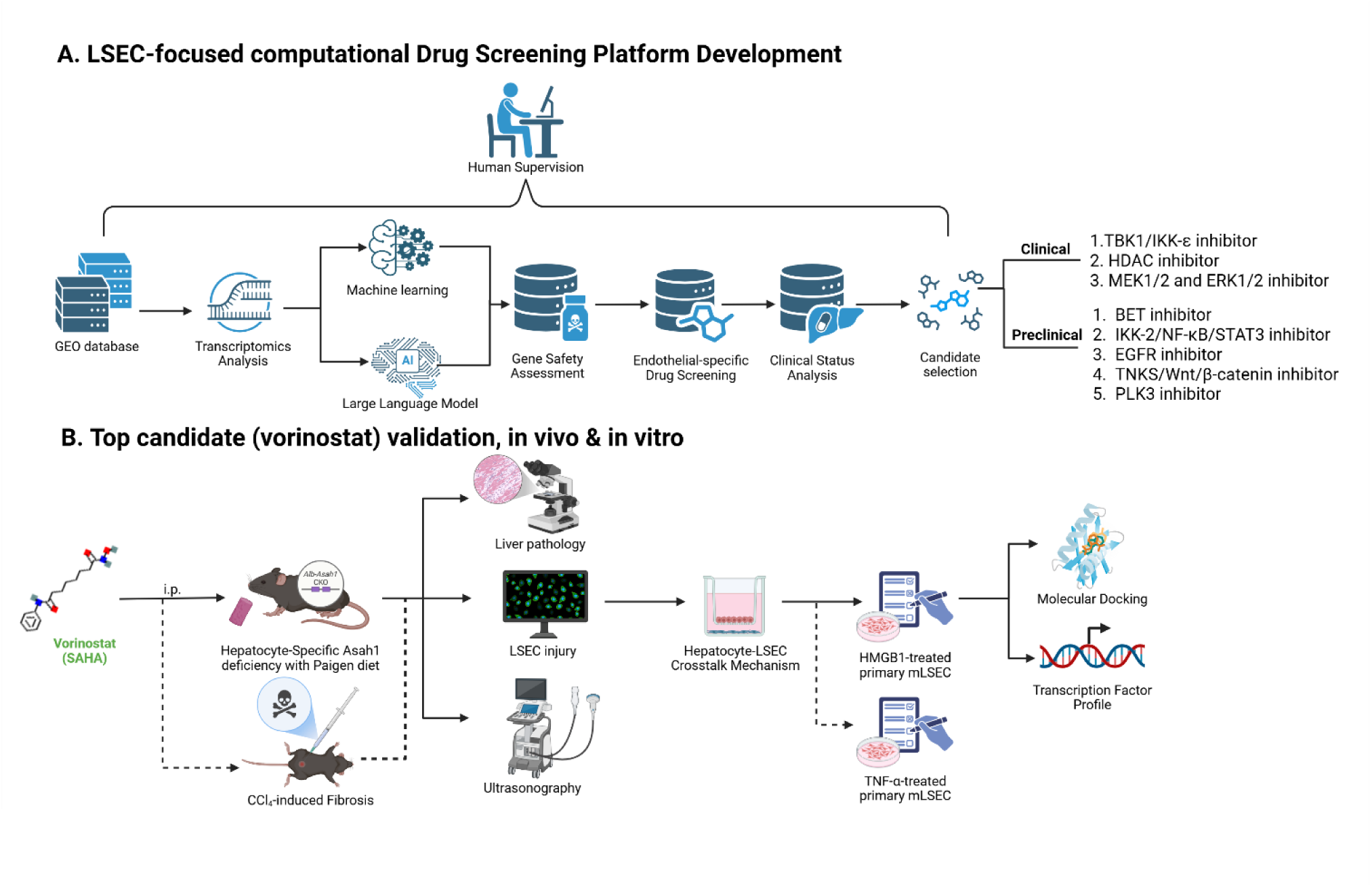

## INTRODUCTION

Liver fibrosis is a common pathological consequence of chronic liver injuries from diverse etiologies, including nonalcoholic fatty liver disease (NAFLD, recently renamed metabolic dysfunction-associated steatotic liver disease [MASLD]) and its progressive form nonalcoholic steatohepatitis (NASH, renamed MASH), viral hepatitis, alcohol-associated, cholestatic, and autoimmune liver diseases[1]. Progressive fibrosis can lead to cirrhosis, liver failure, hepatocellular carcinoma, and even death. Approximately 2 million deaths annually (4% of global deaths) are attributed to liver diseases[2], with advanced fibrosis affecting an estimated 3.3% of the population worldwide, representing a major global health burden[3]. Although resmetirom (2024) and semaglutide (2025) have recently been approved for NASH with moderate-to-advanced fibrosis (F2-F3), both agents target metabolic and inflammatory pathways (thyroid hormone receptor-β and glucagon-like peptide-1 (GLP-1) receptor, respectively) rather than fibrogenic mechanisms [4, 5]. This restricts treatment to noncirrhotic NASH only and does not address fibrosis arising from other etiologies. Thus, there remains a critical unmet need for therapeutic strategies that directly target the fibrogenic process across the spectrum of chronic liver diseases.

Current anti-fibrotic research has predominantly focused on hepatic stellate cells (HSCs), the principal effector cells responsible for extracellular matrix deposition in the fibrotic liver[6, 7]. However, despite decades of HSC-targeted drug development, no HSC-directed therapy has been approved, suggesting that targeting HSCs alone may be insufficient [8, 9]. Liver sinusoidal endothelial cells (LSECs), the most abundant non-parenchymal liver cell population lining the liver sinusoids, have emerged as an important but underestimated cell type during fibrosis pathogenesis [9–11].

Under physiological conditions, LSECs maintain hepatic homeostasis by preserving HSC quiescence through Krüppel-like factor 2 (KLF2)-endothelial nitric oxide synthase (eNOS)-nitric oxide (NO) pathway, regulating immune surveillance, inflammatory responses, vasodilation, and facilitating material exchange through their unique fenestrated structure [12, 13]. These features are critical for the delivery of nutrients, metabolites, oxygen, and therapeutic agents to liver parenchymal cells and HSCs. During chronic liver injury, LSECs undergo capillarization (loss of fenestrae and formation of a basement membrane), which precedes HSC activation. LSEC dysfunction represents an early and potentially reversible event in fibrogenesis, with capillarization observed as early as one week of choline-deficient, L-amino acid-defined (CDAA) diet treatment, preceding the onset of inflammation and fibrosis [14]. Importantly, restoration of LSEC differentiation has been shown to revert activated HSCs to quiescence and promote fibrosis regression[15]. A recent study showed that pharmacological targeting of LSEC angiocrine signaling can treat liver fibrosis in preclinical models and early clinical studies[16]. These findings position LSECs not merely as passive bystanders but as an upstream regulatory cell type whose functional state governs HSC activation and fibrosis resolution. Despite growing recognition of LSECs as critical regulators of fibrosis initiation and progression, systematic LSEC-targeted drug discovery remains largely unexplored.

Connectivity Map (CMap)-based drug repurposing has shown promise in identifying therapeutic candidates by matching disease gene signatures with drug-induced transcriptomic profiles[17]. However, existing approaches typically lack cell-type specificity and thus do not account for the unique biology of LSECs. Furthermore, many CMap-based studies still rely on threshold-defined disease signatures, which may not prioritize disease-relevant, cell-type-specific, or safely targetable genes. To address these gaps, we developed an LSEC-focused computational drug screening platform that integrates LSEC-specific transcriptomic analysis across fibrotic liver disease stages, tiered gene signature selection combining machine learning (ML)-based stability selection, large language model (LLM)-assisted literature curation, comprehensive gene safety assessment, and CMap-based drug screening using human endothelial-specific transcriptomic profiles.

Using this platform, we identified 6 clinical-stage and 8 pre-clinical drug candidates with LSEC-protective potential spanning multiple target classes. Among these, vorinostat (SAHA), a US Food and Drug Administration (FDA)-approved pan-histone deacetylase (HDAC) inhibitor, was selected for experimental validation in two independent mouse models of liver fibrosis. We also investigated the hepatocyte-to-LSEC paracrine injury mechanism and the potential intracellular mechanism underlying SAHA’s endothelial-protective effect.

## METHODS

### Transcriptomic Data Analysis and Functional Enrichment

All transcriptomic analyses were performed using R (version 4.3.2, https://www.r-project.org). For cross-species consistency, human gene symbols (italicized, uppercase) are used throughout the text unless species-specific context requires otherwise.

For the LSEC-specific data input of the computational platform, three public transcriptomic datasets were obtained from the Gene Expression Omnibus (GEO) database (National Center for Biotechnology Information [NCBI], https://www.ncbi.nlm.nih.gov/geo): GSE166504 (mouse liver single-cell RNA sequencing [scRNA-seq], chow vs. high-fat high-fructose diet [HFHFD] at 15 and 30 weeks)[18]; GSE140994 dataset (primary mouse LSEC bulk RNA sequencing, control vs. CDAA diet at 10 weeks)[19], and GSE136103 (human liver scRNA-seq, healthy vs. cirrhotic patients)[20].

For scRNA-seq datasets, LSEC subpopulations were identified by canonical markers and extracted using Seurat R package (v5.2.1)[21]. For GSE136103, standard quality control (500-6,000 features, <20% mitochondrial content), log-normalization, principal component analysis (PCA), and graph-based clustering were performed. LSEC subpopulations were identified at the cluster level using canonical markers (*STAB2, CLEC4G, PECAM1*), extracted, and further purified by requiring expression of at least one canonical LSEC marker. For GSE166504, endothelial cells from the authors’ annotations were further verified using a panel of 13 positive LSEC markers (*STAB1, STAB2, CLEC4G, CLEC4M, FCGR2B, LYVE1, MRC1, PLVAP, GPR182, OIT3, DNASE1L3, CLEC1B, CLEC14A*) and negative marker panels for contaminating cell types to distinguish LSECs from other endothelial subpopulations. Pseudobulk expression profiles were generated by summing raw counts across all cells within each biological replicate.

Differentially expressed gene (DEG) analysis was performed using edgeR package (v4.0.16)[22] for scRNA-seq pseudobulk datasets, or limma R package (v3.58.1)[23] for the bulk RNA-seq dataset (GSE140994). Gene set variation analysis (GSVA) was performed using the GSVA R package (v1.50.5)[24] across 12 Hallmark pathways (obtained via msigdbr R package, v25.1.1, species-matched)[25], grouped into inflammation, fibrosis, and endothelial dysfunction categories. Gene Ontology (GO) enrichment analysis was performed using clusterProfiler R package (v4.10.1)[26].

For the assessment of SAHA on endothelial transcriptomic profile, publicly available RNA-seq data from SAHA-treated human aortic endothelial cells (HAEC) were obtained from GEO (GSE54912). GSVA was performed across 12 Hallmark pathways as described above. Differentially expressed transcription factors were identified and compared between control and SAHA-treated conditions.

### LSEC-specific Gene Signature Selection

LSEC gene signatures were selected by a tiered strategy. For Tier 1 genes, stability selection[27] using elastic net logistic regression (glmnet R package [28], v4.1.8; α = 0.5, 1,000 bootstrap iterations) was applied to FDR < 0.25 DEGs within each pathway. Input was capped at 30 (GSE166504 and GSE136103) or 50 genes (GSE140994) per pathway. Genes selected in ≥50% of iterations were retained. For Tier 2 genes, the top 10 genes per pathway (excluding Tier 1) were evaluated using Claude Opus 4.0 (Anthropic, https://claude.ai) for evidence of endothelial/liver-protective effects of the proposed intervention. Each gene received a recommendation with supporting PubMed identifiers (PMIDs), followed by mandatory human expert validation. If combined Tier 1 + 2 genes were fewer than 10 (up- or downregulated), Tier 3 supplementary genes were included by p-value ranking and human evaluation.

Selected Tiered gene signatures then underwent safety assessment with three databases: DepMap CRISPR common essentials (Broad Institute, https://depmap.org)[29], Catalogue of Somatic Mutations in Cancer (COSMIC) Cancer Gene Census (v103, https://cancer.sanger.ac.uk/cosmic)[30], and ClinVar variant summary (NCBI, https://www.ncbi.nlm.nih.gov/clinvar)[31]. Further literature review was performed to identify other potential side effects. All the results were finally reviewed and approved manually.

### Computational Drug Screening

Drug screening was performed using the human umbilical vein endothelial cell (HUVEC)-specific data in CMap database (Library of Integrated Network-Based Cellular Signatures [LINCS] L1000 platform, https://clue.io)[32]. Twelve queries (4 comparisons × 3 categories) were submitted, and results were filtered by quality control (QC)_Pass = 1, small molecule compounds only, transcriptional activity score (TAS) ≥ 0.2, and n ≥ 3. No positive connectivity (maximum raw connectivity score ≤ 0.05) was allowed across all categories. Drug candidates were required to show disease-signature reversal activity, defined as raw connectivity score < −0.3 in at least one human dataset category (GSE136103) or at least two mouse dataset categories (GSE166504, GSE140994), then ranked by pathway coverage and mean connectivity score. Finally, drug candidates underwent clinical status analysis (FDA approval status, drug targets, mechanism, administration route, hepatotoxicity risk and other severe side effects) to identify repurposing potential.

### High mobility group box 1 (HMGB1)-LSEC Receptor Interaction Analysis

Mouse and human HMGB1-receptor interaction in hepatocyte-LSEC intercellular communication was quantified using the formulation: interaction strength = mean HMGB1 expression (hepatocytes) × mean receptor expression (LSECs). Interaction dynamics were analyzed across disease stages for mouse (GSE166504: chow, HFHFD 15 weeks, HFHFD 30 weeks) and human (GSE136103: healthy, cirrhotic) datasets. LSEC receptors were classified as pro-inflammatory (Advanced glycosylation end-product specific receptor [AGER, also known as RAGE], Toll-like receptor [TLR]4, TLR2), anti-inflammatory (CD24, thrombomodulin [THBD]), chemotaxis/regeneration (C-X-C motif chemokine receptor 4 [CXCR4]), and secondary (hepatitis A virus cellular receptor 2 [HAVCR2, also known as TIM-3], integrin subunit alpha M [ITGAM], triggering receptor expressed on myeloid cells 1 [TREM1]).

### Molecular Docking

Molecular docking was performed to compare binding of glycyrrhizin (positive control) and SAHA to HMGB1 and its LSEC receptors. Ligand structures were obtained from the PubChem database (https://pubchem.ncbi.nlm.nih.gov): glycyrrhizin (CID: 14982), SAHA (CID: 5311). Two-dimensional structure (CID: 14982) was converted to three-dimensional conformations, and all ligands were energy-minimized using Open Babel (MMFF94 force field) within PyRx (version 0.8, https://pyrx.sourceforge.io)[33]. Protein structures were obtained from the RCSB Protein Data Bank (PDB, https://www.rcsb.org): HMGB1 A+B box (PDB: 2YRQ), thrombin/thrombomodulin (THBD) complex (PDB: 1HLT), TLR4/MD-2 complex (PDB: 3FXI), and CXCR4 (PDB: 3ODU). Receptors were prepared in PyMOL (version 3.0.3, https://pymol.org), including removal of co-crystallized ligands, water molecules, and non-protein heteroatoms.

Docking was performed using AutoDock Vina[34] (integrated with PyRx, exhaustiveness = 8), generating up to ten poses per ligand-receptor pair. Interaction profiling of top-ranked poses was performed using the Protein-Ligand Interaction Profiler[35] (PLIP, https://plip-tool.biotec.tu-dresden.de/plip-web/plip) web server. Three-dimensional docking poses and two-dimensional interaction diagrams were visualized using Discovery Studio Visualizer (BIOVIA, 2019).

### Chemicals and Reagents

SAHA and oleic acid (OA) were purchased from Cayman Chemical (Ann Arbor, MI). Recombinant mouse HMGB1, mouse TNF-α, carbon tetrachloride (CCl_4_), palmitic acid (PA), 7-ketocholesterol (7K), and glycyrrhizin (Gly) were purchased from Sigma-Aldrich (St. Louis, MO). For in vivo treatment, SAHA was dissolved in a vehicle consisting of 10% dimethyl sulfoxide (DMSO), 45% polyethylene glycol 400 (PEG400), and 45% sterile water. CCl_4_ was diluted in corn oil (v/v CCl_4_: corn oil = 1:4). For in vitro treatment, SAHA and Gly were dissolved in DMSO. Recombinant mouse HMGB1 and TNF-α were reconstituted in sterile phosphate-buffered saline (PBS) + 0.1% bovine serum albumin (BSA). PA was dissolved in 0.1 mol/L NaOH at 75°C and diluted in fatty acid-free BSA solution. OA and 7K were dissolved in ethanol.

### Animal Models and In Vivo Treatment

All animal studies were conducted under the National Institutes of Health (NIH) *Guide for the Care and Use of Laboratory Animals*. All animal experiments were approved by the Institutional Animal Care and Use Committee (IACUC) of University of Houston, College of Pharmacy. Mice were housed in a temperature- and humidity-controlled room with a 12-hour light/12-hour dark cycle, provided with rodent chow and water ad libitum. Hepatocyte-specific *Asah1* knockout mice (*Asah1*^fl/fl^/*Alb*^Cre^) were generated by crossing *Asah1* flox mice (*Asah1*^fl/fl^/wild type, WT) with *Alb*-Cre mice, and genotyped as previously described[36]. WT C57BL/6J mice were bred and maintained at the University of Houston animal facility.

#### Hepatocyte-specific Asah1 knockout mice treatment

Male *Asah1*^fl/fl^/WT and *Asah1*^fl/fl^/*Alb*^Cre^ mice (C57BL/6J background; aged 8-20 weeks) were fed a high-fat, high-cholesterol Paigen diet (PD, Research Diets D12336) from week 0. SAHA (15 mg/kg/day, intraperitoneally [i.p.], 5 times/week) or vehicle (10% DMSO, 45% PEG400, 45% sterile water) was administered from week 6 to week 20. Portal venous hemodynamics and cardiac function were assessed by ultrasonography at weeks 12 and 20. Mice were sacrificed at week 20 after overnight fasting. Livers and plasma were collected for downstream analysis. For HMGB1 quantification **(Figure 5B)**, hepatic and plasma samples from *Asah1*^fl/fl^/WT and *Asah1*^fl/fl^/*Alb*^Cre^ mice fed normal diet (ND) or PD for 20 weeks were obtained from our previous study[36].

#### CCl_4_-induced liver fibrosis model treatment

Male WT C57BL/6J mice (aged 8-12 weeks) received CCl_4_ (v/v CCl_4_: corn oil = 1:4, 5 mL/kg, i.p., twice weekly) or vehicle (corn oil) from week 0. SAHA (15 mg/kg/day, i.p., 5 times/week) or vehicle was administered from week 1. Portal venous hemodynamics were assessed by ultrasonography at week 4. Mice were sacrificed at week 4 after overnight fasting.

### Cell Culture and In Vitro Treatments

Primary mouse liver sinusoidal endothelial cells (LSECs, C57BL/6 background, Cat. No. C57-6017; Cell Biologics, Chicago, IL) were cultured in Complete Endothelial Cell Medium (Cat. No. M1168; Cell Biologics) and used within passages 3-6, according to the manufacturer’s instruction. Mouse macrophage cell line J774 was purchased from American Type Culture Collection (ATCC, Manassas, VA) and maintained in Dulbecco’s Modified Eagle Medium (DMEM) (high glucose) supplemented with 10% fetal bovine serum (FBS). Human hepatocellular carcinoma cell line HepG2 was purchased from ATCC and maintained in Eagle’s minimum essential medium (EMEM) with 10% FBS. All cells were cultured in a humidified incubator with 5% CO_2_ at 37°C.

For the assessment of SAHA on HMGB1-induced injury, primary LSECs were treated with recombinant HMGB1 (1 μg/mL) with or without SAHA (0, 2.5, 5, or 10 μM) for 48 hours. For the assessment of SAHA on TNF-α-induced injury, primary LSECs were treated with TNF-α (40 ng/mL) with or without SAHA (0 or 10 μM) for 12 hours. For KLF2 validation, primary LSECs were treated with SAHA (0, 2.5, 5, or 10 μM) for 24 hours. For HMGB1 quantification, HepG2 cells were transfected with human *ASAH1* small interfering RNA (siRNA) or negative control siRNA (siNC) (Santa Cruz Biotechnology, Dallas, TX) using silentFect Lipid Reagent (Bio-Rad, Hercules, CA) for 48 hours, followed by treatment with vehicle or a lipid mixture consisting of free fatty acids (FA, 300 μM; OA: PA = 2:1) and 7K (40 μM) for 24 hours. Cell lysate and culture supernatant were collected for enzyme-linked immunosorbent assay (ELISA) quantification. For Transwell co-culture, HepG2 hepatocytes were cultured in the upper chamber of Transwell inserts and transfected with siNC or si*ASAH1* for 48 hours. Primary LSECs were cultured separately in the lower chamber. After transfection, co-culture was initiated with FA + 7K treatment in the upper chamber and vehicle or Gly (60 μM) in the lower chamber for 24 hours. LSEC gene expression was assessed by qPCR.

### Ultrasonography

Mouse portal venous hemodynamics and cardiac function were assessed using a Vevo 3100 Imaging System (VisualSonics, Toronto, ON, Canada). Portal venous hemodynamics were analyzed using B-mode, color Doppler mode, and pulsed wave Doppler mode, as described before[36]. Maximum velocity (V_max_), minimum velocity (V_min_), and mean velocity (V_mean_) were analyzed using Vevo LAB (v2.1.0, VisualSonics). Pulsatility index (PI) was calculated as PI = (V_max_ - V_min_)/V_max_. Cardiac function assessment was performed by M-mode echocardiography, as previously described[37]. Ejection fraction (EF), stroke volume, left ventricular internal diameter in diastole (LVID;d) and systole (LVID;s), and left ventricular volume in diastole (LV Vol;d) and systole (LV Vol;s) were analyzed using Vevo LAB (v2.1.0).

### Biochemical Assays and ELISA

Plasma alanine aminotransferase (ALT) and aspartate aminotransferase (AST) activities, and total cholesterol and triglyceride concentrations in mouse plasma and liver tissue were determined by commercial kits according to the manufacturer’s protocols (BioAssay Systems, Hayward, CA). HMGB1 concentrations of mouse liver tissue, mouse plasma, cell lysate, and cell culture supernatant were measured by ELISA (Elabscience, Houston, TX) according to the manufacturer’s protocol. Hepatic lipid concentrations and HMGB1 levels were normalized to total protein, quantified by bicinchoninic acid (BCA) assay (Thermo Fisher, Waltham, MA).

### Pathological Staining

Liver paraffin sections (4-μm-thick) or frozen liver sections (8-μm-thick) were used for hematoxylin and eosin (H&E, Teomics, Houston, TX) staining, Sirius Red (Sigma-Aldrich) staining, or terminal deoxynucleotidyl transferase-mediated dUTP nick-end labeling (TUNEL, Roche, Basel, Switzerland) staining according to the manufacturer’s protocol.

### Immunofluorescence Staining

Immunofluorescence (IF) staining was performed as previously described[36]. Briefly, for hepatic tissue staining, frozen liver sections (8-μm-thick) were fixed with acetone, washed with PBS, and blocked with 10% donkey serum for 1 hour. For cell immunofluorescence staining, primary LSECs were fixed with 4% paraformaldehyde, washed with PBS, and blocked with 5% donkey serum for 30 minutes. Sections or cells were then incubated with primary antibodies at room temperature for 1 hour and overnight at 4°C, followed by incubation with Alexa Fluor 488- and/or Alexa Fluor 555-labeled secondary antibodies (Thermo Fisher) for 1 hour at room temperature. Slides were mounted with Vectashield PLUS antifade mounting medium with 4’,6-diamidino-2-phenylindole (DAPI, Vector Laboratories, Newark, CA) and imaged using an IX73 imaging system (Olympus, Tokyo, Japan). Primary CD45, CD11b, Collagen I, intercellular adhesion molecule 1 (ICAM-1), vascular cell adhesion molecule 1 (VCAM-1), and KLF2 antibodies were purchased from Abcam (Cambridge, UK). C-C motif chemokine ligand 2 (CCL2) primary antibody was purchased from Millipore (Burlington, MA). CD34 primary antibody was purchased from Cell Signaling Technology (Danvers, MA). Endothelin-1 (ET-1) primary antibody was purchased from Invitrogen (Carlsbad, CA). For liver co-immunofluorescence staining, IB4 (isolectin B4; Invitrogen) was used as an endothelial marker in combination with ICAM-1, VCAM-1, CD34, ET-1, or KLF2 antibodies.

### Quantitative polymerase chain reaction (qPCR)

QPCR analysis was performed as previously described[36]. Briefly, total RNA was extracted using TRIzol reagent (Invitrogen) and reverse transcribed using iScript Reverse Transcription Supermix (Bio-Rad, Hercules, CA), according to the manufacturer’s protocol. Gene expression was determined using iTaq Universal SYBR Green Supermix (Bio-Rad). Primers were purchased from Eurofins Scientific (Luxembourg City, Luxembourg), and sequences are listed in **Supplementary Table 1**. For hepatic gene expression analysis, results were normalized to peptidylprolyl isomerase A (*PPIA*). For cell experiments, gene expression was normalized to hypoxanthine phosphoribosyl transferase 1 (*HPRT1*).

### Western Blot

Western blot (WB) analysis was performed as previously described[36]. Briefly, primary LSECs were lysed, boiled at 95°C for 5 minutes, and separated on sodium dodecyl sulfate-polyacrylamide gel electrophoresis (SDS-PAGE). Proteins were transferred to polyvinylidene difluoride membranes, blocked with 5% BSA, and incubated with primary antibodies for KLF2 (Abcam) and GAPDH (Santa Cruz Biotechnology) overnight at 4°C. Bands were incubated with corresponding secondary antibodies (Thermo Fisher) and detected using a LI-COR Western Blot Imaging system (LI-COR, Lincoln, NE). Band intensity was analyzed using ImageJ software (NIH, Bethesda, MD).

### Monocyte Adhesion Assay

Monocyte adhesion assay was performed as previously described[38]. Briefly, J774 mouse macrophage cells were stained with 2 μM Calcein-AM (Cat. No. C3100MP; Thermo Fisher) at 37°C for 10 minutes, seeded onto primary LSECs (treated with TNF-α, with or without SAHA) for 30 minutes. Unbound monocytes were discarded, and LSEC layers with attached monocytes were washed with PBS, fixed with 4% paraformaldehyde for 10 minutes, stained with DAPI for 15 minutes, and imaged using an IX73 imaging system (Olympus).

### Statistical Analysis

Data normality was assessed by Shapiro-Wilk test. For normally distributed data, ordinary one-way analysis of variance (ANOVA) followed by Fisher’s least significant difference (LSD) post hoc test was used. For non-normally distributed data, Kruskal-Wallis test followed by Dunn’s multiple comparisons test was used. Data are presented as mean ± standard error of the mean (SEM). All statistical analyses were performed using GraphPad Prism (version 8.0; GraphPad Software, San Diego, CA). *P* < 0.05 was considered statistically significant. \**P* < 0.05, \*\**P* < 0.01.

### Data Visualization

Heatmaps were generated using pheatmap (v1.0.13) and ComplexHeatmap (v2.18.0) R packages with the viridis color palette. Venn diagrams were generated using ggVennDiagram R package (v1.5.7). CMap connectivity score heatmaps, HMGB1 interaction dot plots, and correlation plots were generated using ggplot2 (v4.0.1) with ggrepel. Chord diagrams were generated using circlize R package (v0.4.17). Protein-protein interaction (PPI) network analysis and visualization were performed using the Search Tool for the Retrieval of Interacting Genes/Proteins (STRING) database (https://string-db.org/)[39].

## RESULTS

### Development of an LSEC-focused computational drug screening platform

To systematically identify drugs that improve LSEC dysfunction (manifested by reversed gene signatures) across fibrotic liver disease progression, we developed a computational screening platform **(Figure 1A)**. Three public datasets from different stages of fibrosis were integrated, including simple steatosis NAFLD stage (GSE166504, chow vs HFHFD 15 weeks), fibrotic NASH stage (GSE166504, chow vs HFHFD 30 weeks, and GSE140994, control vs CDAA diet at 10 weeks), and cirrhosis (GSE136103, healthy vs cirrhotic patients) **(Figure 1A, step 1)**.

**Figure 1.**
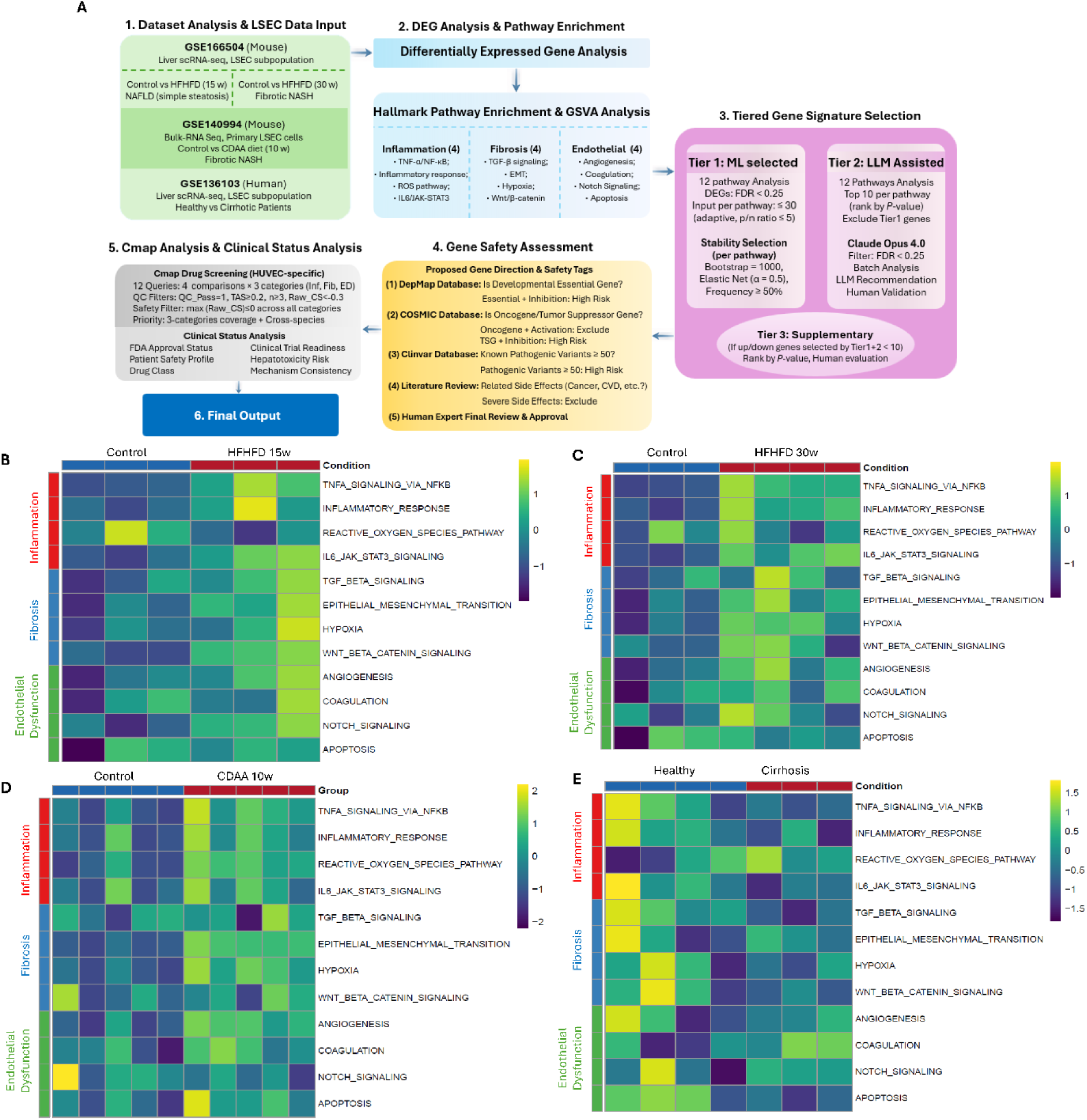
Development of an LSEC-focused computational drug screening platform. **(A)** Platform overview. (1) LSEC transcriptomic data from three public GEO datasets: GSE166504 (mouse liver scRNA-seq; chow vs. HFHFD at 15 and 30 weeks), GSE140994 (mouse primary LSEC bulk RNA-seq; control vs. CDAA diet at 10 weeks), and GSE136103 (human liver scRNA-seq; healthy vs. cirrhosis patients). For scRNA-seq datasets, LSEC subpopulations were identified by canonical markers, extracted, and pseudobulk-aggregated. (2) DEG analysis across four comparisons, followed by Hallmark pathway enrichment and GSVA across 12 pathways grouped into three functional categories: Inflammation, Fibrosis, and Endothelial dysfunction. (3) Tiered gene signature selection combining ML-based stability selection (Tier 1), LLM-assisted curation with human validation (Tier 2), and supplementary genes by *p*-value ranking and human evaluation (Tier 3). (4) Gene safety assessment using DepMap, COSMIC, and ClinVar databases, literature review, and human expert approval. (5) CMap drug screening (HUVEC cell line, 12 queries) with quality control filtering, safety filtering, and clinical status analysis. (6) Final output of prioritized drug candidates. **(B-E)** GSVA heatmaps of 12 Hallmark pathways (Inflammation, Fibrosis, Endothelial Dysfunction) across four comparisons: (B) NAFLD with simple steatosis (mouse, HFHFD 15 weeks, GSE166504) (C) NASH with fibrosis (mouse, HFHFD 30 weeks, GSE166504) (D) NASH with fibrosis (mouse, CDAA 10 weeks, GSE140994) and (E) cirrhosis (human, GSE136103). Abbreviations: GEO, Gene Expression Omnibus; scRNA-seq, single-cell RNA sequencing; HFHFD, high-fat high-fructose diet; CDAA, choline-deficient L-amino acid-defined; NAFLD, nonalcoholic fatty liver diseases; NASH, nonalcoholic steatohepatitis; DEG, differentially expressed gene; GSVA, gene set variation analysis; ROS, reactive oxygen species; EMT, epithelial-mesenchymal transition; ML, machine learning; LLM, large language model; FDR, false discovery rate; DepMap, Dependency Map; COSMIC, Catalogue of Somatic Mutations in Cancer; TSG, tumor suppressor gene; CVD, cardiovascular diseases; CMap, Connectivity Map; HUVEC, human umbilical vein endothelial cell; Inf, inflammation; Fib, fibrosis; ED, endothelial dysfunction; QC, quality control; TAS, transcriptional activity score; CS, connectivity score.

Next, DEG analysis and Hallmark pathway enrichment were performed. Twelve Hallmark pathways were selected and grouped into three categories due to their close relationship with LSEC dysfunction, including inflammation (Tumor necrosis factor-alpha [TNF-α]/nuclear factor kappa B [NF-κB], Inflammatory response, Reactive oxygen species [ROS] pathway, and Interleukin [IL]-6/Janus kinase [JAK]-signal transducer and activator of transcription [STAT]3), fibrosis (Transforming growth factor-beta [TGF-β] signaling, Epithelial-mesenchymal-transition [EMT], Hypoxia, and Wnt/β-catenin), and endothelial dysfunction (Angiogenesis, Coagulation, Notch Signaling, and Apoptosis) **(Figure 1A, step 2)**. GSVA results of these functional categories are shown in **Figure 1B-E**.

GSVA analysis revealed progressive activation of inflammation- and fibrosis-related pathways from simple steatosis to fibrotic NASH in mouse LSECs **(Figure 1B-D)**. TNF-α/NF-κB and EMT signaling pathways were significantly upregulated across both stages, while, the Inflammatory response, IL6/JAK-STAT3, ROS, Hypoxia, and Angiogenesis pathways reached significance primarily at the fibrotic NASH stage. EMT was consistently enriched across all three mouse comparisons. TGF-β signaling showed a modest increasing trend without reaching statistical significance. In contrast, the human cirrhosis dataset (GSE136103) exhibited a distinct pattern **(Figure 1E)**. TNF-α/NF-κB, IL6/JAK-STAT3, and TGF-β showed negative enrichment in cirrhotic patients relative to healthy LSECs.

### Identification of disease-stage-specific LSEC gene signatures

To identify LSEC-specific gene signatures across different fibrotic stages, we employed a computational tiered strategy **(Figure 1A, step 3).** For Tier 1 genes, machine learning-based stability selection using elastic net regression was applied to prioritize repeatedly selected genes under bootstrap resampling (Frequency ≥ 50%). For remaining top-ranked DEGs, large language model (Claude Opus 4.0) was employed to assist literature curation, followed by mandatory human validation (Tier 2). If Tier 1 + Tier 2 combined yielded fewer than 10 up- or downregulated genes per comparison, supplementary genes (Tier 3) were added by *p*-value ranking and human evaluation. All selected genes then underwent safety assessment using DepMap (to identify developmental essential genes), COSMIC (to identify oncogenes and tumor suppressor genes), and ClinVar (to identify numbers of known clinical pathological variants) databases, followed by literature review (to identify other potential side effects such as cancer and cardiovascular diseases), and final human expert review and approval **(Figure 1A, step 4).**

The resulting gene signatures across the four comparisons are shown in **Figure 2A-D**. The number of selected genes varied by dataset and disease stages. Venn diagram analysis of upregulated signatures **(Figure 2E)** showed that most genes were dataset-specific or shared between two to three comparisons. A similar pattern was observed for downregulated signatures **(Figure 2F)**. A consensus heatmap of genes identified in at least two datasets **(Figure 2G)** revealed a core set of consistently dysregulated LSEC genes spanning disease stages and species (mouse and human). Upregulated consensus genes included pro-inflammatory mediators (*CCL2, C-X-C motif chemokine ligand 2 [CXCL2], IL18, IL1B*), adhesion molecules (*VCAM-1, CD44*), and extracellular matrix regulators (*matrix metalloproteinase 14 [MMP14], S100A4*). Downregulated consensus genes included endothelial-protective transcription factors (*KLF2, KLF10*), and LSEC differentiation markers (*stabilin 1 [STAB1], scavenger receptor class F member 1 [SCARF1]*).

**Figure 2.**
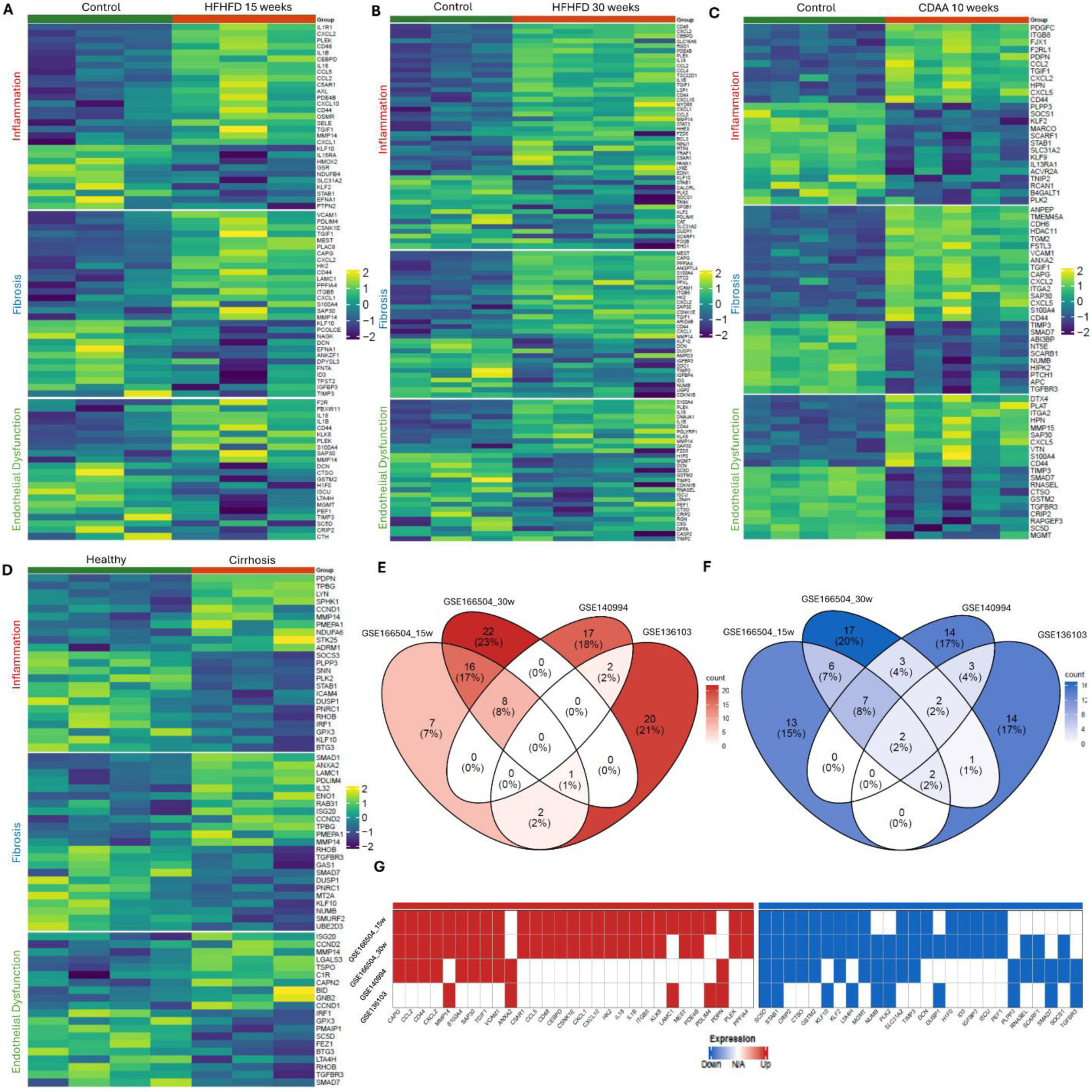
LSEC-specific gene signatures across fibrotic liver disease progression (A-D) LSEC-specific gene signature heatmaps organized by three functional categories for each comparison: (A) chow vs. HFHFD 15 weeks (simple steatosis) and (B) HFHFD 30 weeks (fibrotic NASH) from GSE166504; (C) control vs. CDAA 10 weeks (fibrotic NASH) from GSE140994; (D) healthy vs. cirrhotic patients from GSE136103. **(E-F)** Venn diagrams showing overlap of upregulated (E) and downregulated (F) tiered gene signatures across four comparisons. **(G)** Heatmap of consensus genes identified in at least two datasets. Red indicates upregulation, blue indicates downregulation, and white indicates the gene was not selected as a gene signature in that dataset. Abbreviations: HFHFD, high-fat high-fructose diet; CDAA, choline-deficient L-amino acid-defined; NASH, nonalcoholic steatohepatitis.

### CMap drug screening identifies clinical and pre-clinical candidates with LSEC-protective potential across fibrotic liver disease stages

Next, gene signatures that passed safety assessment were queried against the CMap database across 12 queries (4 comparisons × 3 functional categories). Since LSEC-specific perturbational profiles are not available in CMap, HUVEC profiles were used as a pragmatic endothelial perturbation resource. Therefore, CMap results were interpreted as a prioritization layer requiring downstream validation in primary LSECs and in vivo fibrosis models. Afterwards, the candidate drugs with HUVEC-specific data were filtered by quality control criteria and a safety filter, as described in **Methods**. The resulting connectivity score heatmap of the top 30 drug candidates is shown in **Figure 3A**. The number of reversed functional categories across datasets, reflecting the breadth of cross-stage signature reversal for each candidate, is shown in **Figure 3B**.

**Figure 3.**
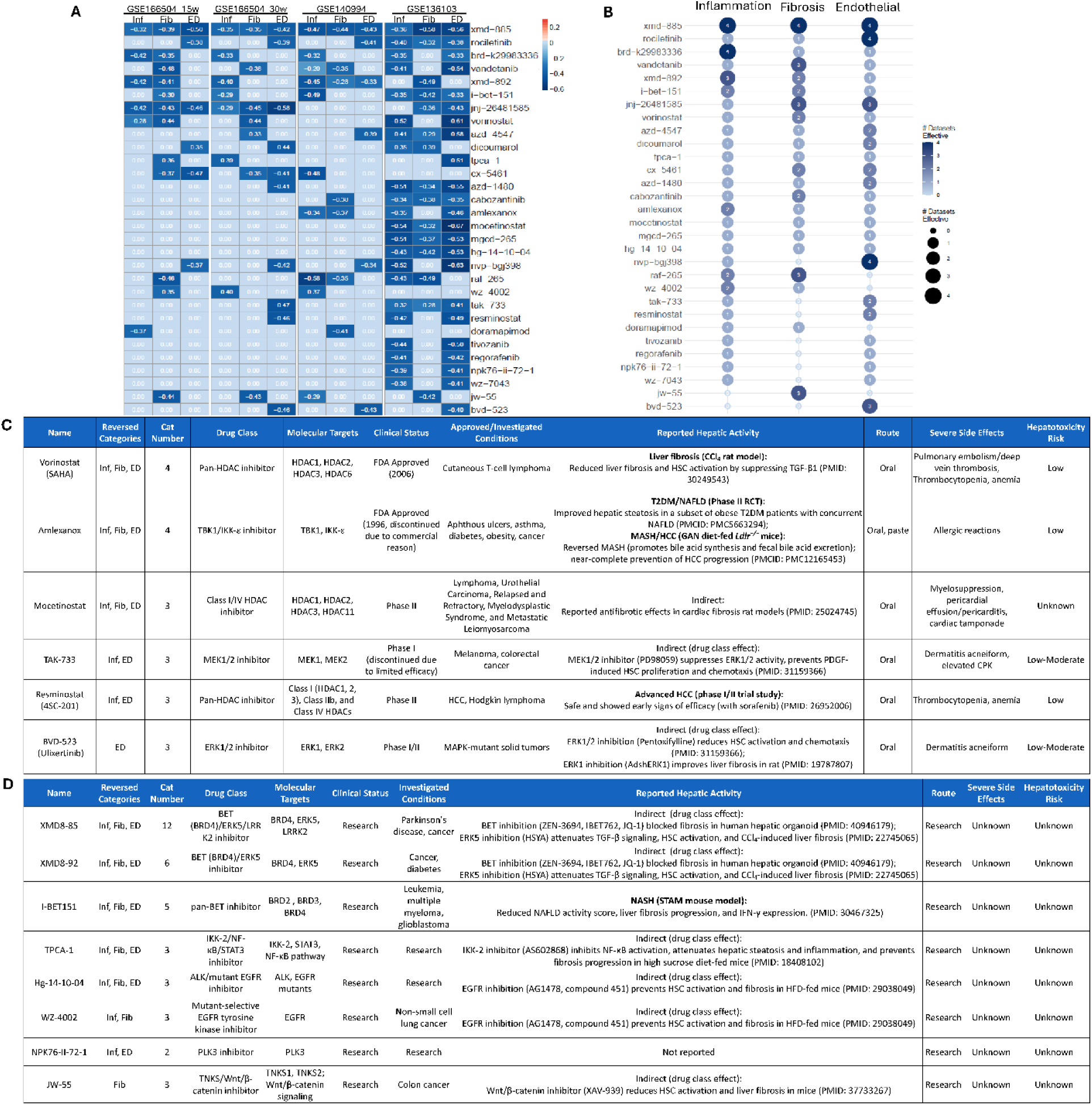
CMap analysis identifies clinical and preclinical candidates across fibrotic progression with LSEC-protective potential. **(A)** CMap connectivity score heatmap for top 30 drug candidates, obtained across 12 queries (4 comparisons × 3 functional categories). Negative connectivity scores indicate reversal of the disease-associated LSEC gene signatures. Darker blue indicates stronger effect. **(B)** Bubble plot showing the number of reversed functional categories per drug across 3 stages. **(C)** Clinical stage and **(D)** Preclinical drug candidates with drug class, targets, clinical status, and safety profile. SAHA was selected for proof-of-concept validation based on clinical availability, prior anti-fibrotic evidence, and strong LSEC signature-reversal activity. Abbreviations: Inf, inflammation; Fib, fibrosis; ED, endothelial dysfunction; Cat, category; HDAC, histone deacetylase; TBK1, TANK-binding kinase 1; IKK, inhibitor of nuclear factor kappa-B kinase; MEK, mitogen-activated protein kinase kinase; ERK, extracellular signal-regulated kinase; BET, bromodomain and extra-terminal domain; BRD, bromodomain-containing protein; LRRK2, leucine-rich repeat kinase 2; STAT3, signal transducer and activator of transcription 3; NF-κB, nuclear factor kappa-light-chain-enhancer of activated B cells; ALK, anaplastic lymphoma kinase; EGFR, epidermal growth factor receptor; PLK3, polo-like kinase 3; TNKS, tankyrase; CCl₄, carbon tetrachloride; HSC, hepatic stellate cells; TGF-β, transforming growth factor-beta; T2DM, type 2 diabetes mellitus; RCT, randomized controlled trial; MASH, metabolic dysfunction-associated steatohepatitis; NAFLD, nonalcoholic fatty liver disease; NASH, nonalcoholic steatohepatitis; GAN, Gubra-Amylin NASH; LDLR, low-density lipoprotein receptor; HCC, hepatocellular carcinoma; STAM, stelic Animal Model; IFN-γ, interferon gamma; CPK, creatine phosphokinase.

To prioritize candidates with clinical repurposing potential, we performed clinical status analysis evaluating FDA approval status, hepatotoxicity risk, drug class, and mechanism of action. Clinical drug candidates and detailed information are shown in **Figure 3C**, including HDAC inhibitors (vorinostat, mocetinostat, resminostat), TANK-binding kinase 1 (TBK1)/ inhibitor of nuclear factor kappa-B kinase (IKK)-ε inhibitor (Amlexanox), mitogen-activated protein kinase kinase (MEK)1/2 inhibitor (TAK-733), and extracellular signal-regulated kinase (ERK1/2) inhibitor (BVD-523). Preclinical candidates are shown in **Figure 3D**, including bromodomain and extra-terminal domain (BET) inhibitor (XMD8-85, XMD8-92, I-BET151), IKK-2/NF-κB/STAT3 inhibitor (TPCA-1), epidermal growth factor receptor (EGFR) inhibitors (Hg-14-10-04, WZ-4002), polo-like kinase 3 (PLK3) inhibitor (NPK76-II-72-1), and tankyrase/Wnt/β-catenin inhibitor (JW-55).

Importantly, an independent Phase II clinical trial of Amlexanox has demonstrated improved hepatic steatosis in a subset of obese type 2 diabetes mellitus (T2DM) patients with concurrent NAFLD [40]. Preclinical evidence also demonstrated that Amlexanox reversed MASH and prevented hepatocellular carcinoma (HCC) in Gubra-Amylin NASH (GAN) diet-fed *Ldlr*^−/−^ mice[41]. Beyond Amlexanox, the anti-fibrosis effect of SAHA by inhibiting HSC activation has been demonstrated in another independent research using CCl_4_ rat model[42]. Resminostat has been evaluated in advanced HCC (Phase I/II trial)[43]. The preclinical candidate, I-BET151, has been shown to reduce NAFLD activity score and liver fibrosis progression in a NASH model using stelic animal model (STAM) mice[44].

Indirect anti-fibrotic evidence (reported drug class effects) include MEK1/2 and/or ERK1/2 inhibition[45, 46] (TAK-733, BVD-523), BET/ERK5 inhibition[47, 48] (XMD8-85, XMD8-92), IKK-2 inhibition[49] (TPCA-1), EGFR inhibition[50] (Hg-14-10-04, WZ-4002), and Wnt/β-catenin inhibition[51] (JW-55). In this study, SAHA was selected for proof-of-concept validation based on FDA approval status, established formulation, prior anti-fibrotic evidence, and strong LSEC signature-reversal activity in our screen [42].

### Vorinostat attenuates hepatic injury, LSEC dysfunction, and portal hemodynamic abnormalities in hepatocyte-specific *Asah1*-deficient PD mice

We recently reported that hepatocyte-specific deficiency of *Asah1* (encoding acid ceramidase and mediating ceramide degradation) disrupts hepatic lipid homeostasis and promotes fibrotic NASH progression in mice fed a PD[36]. To validate the anti-fibrotic and LSEC-protective effect of SAHA in vivo, we first employed this recently established model, in which disrupted sphingolipid metabolism drives progressive NASH with inflammation, fibrosis, and hepatocyte-to-non-parenchymal cell injury. This model also develops portal hemodynamic abnormalities and systolic cardiac dysfunction, enabling assessment of whether SAHA’s therapeutic effect extends beyond hepatic injury. The experimental design is shown in **Figure 4A**.

**Figure 4.**
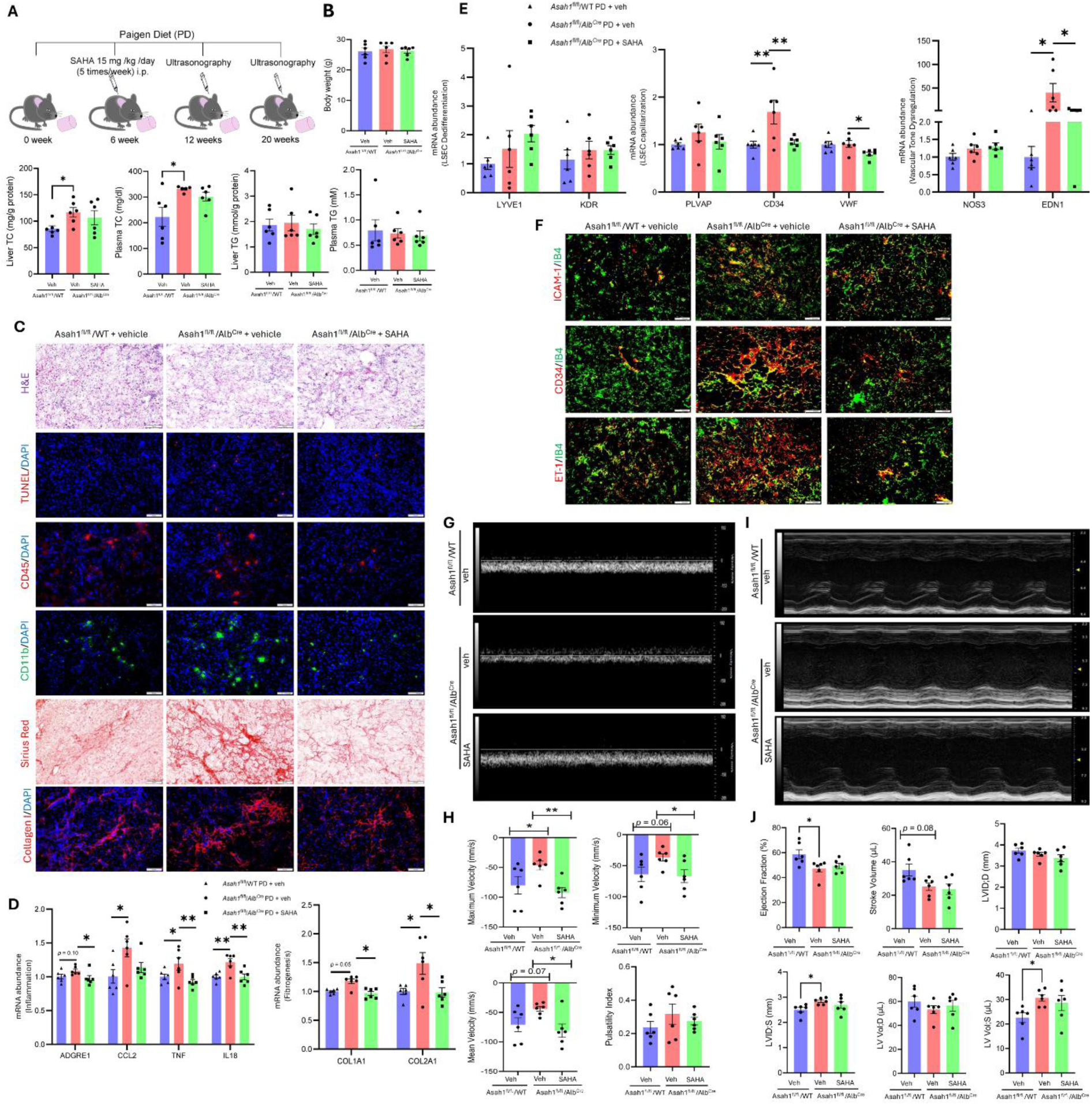
SAHA improves hepatic injury, LSEC dysfunction, and portal hemodynamics in hepatocyte-specific *Asah1*-deficient mice. **(A)** Experimental design. *Asah1*^fl/fl^/*Alb*^Cre^ mice (hepatocyte-specific deletion of *Asah1*) and *Asah1* floxed (*Asah1*^fl/fl^/WT) mice were fed with PD from week 0, treated with vehicle or SAHA (15 mg/kg/day, i.p.) from week 6, and sacrificed at week 20. Portal venous hemodynamics and cardiac function were assessed by ultrasonography at weeks 12 and 20. **(B)** Mouse body weight (g), hepatic TC (mg/g protein), plasma TC (mg/dL), hepatic TG (mmol/g protein), and plasma TG (mM). Hepatic lipid concentrations were normalized to total protein by BCA assay. **(C)** Frozen liver sections (8-μm-thick) were used for pathological staining (H&E, Sirius Red) or immunofluorescent staining (TUNEL, CD45, CD11b, or Collagen I, with DAPI). **(D)** Hepatic gene expression of inflammation and fibrosis markers were quantified by qPCR, normalized to *PPIA*. **(E)** Hepatic gene expression of LSEC related markers (LSEC Dedifferentiation, Capillarization, and Vascular tone dysregulation) were quantified by qPCR, normalized to *PPIA*. **(F)** Frozen liver sections (8-μm-thick) were used for co-immunofluorescence staining, including ICAM-1, CD34, or ET-1, with IB4 (endothelial marker). **(G-H)** Representative portal vein Doppler waveforms (week 20) and hemodynamic quantification including maximum velocity (mm/s), minimum velocity (mm/s), mean velocity (mm/s), and pulsatility index. **(I-J)** Representative echocardiography images (week 20) and cardiac function parameters including Ejection Fraction (%), Stroke Volume (μL), LVID D (mm), LVID S (mm), LV Vol D (μL), and LV Vol S (μL). Mean ± SEM; n = 5-6/group. \**p* < 0.05, \*\**p* < 0.01. Abbreviations: i.p., intraperitoneal; TC, total cholesterol; TG, triglyceride; BCA, bicinchoninic acid; PPIA, peptidylprolyl isomerase A; H&E, hematoxylin and eosin; TUNEL, terminal deoxynucleotidyl transferase-mediated dUTP nick-end labeling; DAPI, 4’,6-diamidino-2-phenylindole; ADGRE1, adhesion G protein-coupled receptor E1 (also known as F4/80); CCL2, C-C motif chemokine ligand 2; TNF, tumor necrosis factor; IL18, interleukin-18; COL1A1, collagen type I alpha 1 chain; COL2A1, collagen type II alpha 1 chain; LYVE1, lymphatic vessel endothelial hyaluronan receptor 1; KDR, kinase insert domain receptor (encoding VEGFR2); PLVAP, plasmalemma vesicle-associated protein; CD34, cluster of differentiation 34; VWF, von Willebrand factor; NOS3, nitric oxide synthase 3; EDN1, endothelin 1; IB4, isolectin B4; ICAM-1, intercellular adhesion molecule 1; ET-1, endothelin-1; LVID D, left ventricular internal diameter in diastole; LVID S, left ventricular internal diameter in systole; LV Vol D, left ventricular volume in diastole; LV Vol S, left ventricular volume in systole.

SAHA treatment did not affect body weight and lipid concentrations (including hepatic total cholesterol, plasma total cholesterol, hepatic triglyceride, and plasma triglyceride) **(Figure 4B)**. Besides, H&E staining showed the minimal effect of SAHA on hepatic steatosis **(Figure 4C).** However, cell death (TUNEL staining), immune cell infiltration (CD45-positive immune cells and CD11b-positive myeloid cells), and collagen deposition (red area of Sirius Red staining and Collagen I staining), were all reduced by SAHA **(Figure 4C)**. The gene expression levels of hepatic inflammation and fibrosis markers were then evaluated by qPCR. SAHA significantly reduced inflammation-related genes including *ADGRE1*, *TNF* and *IL18*, while *CCL2* showed a decreasing trend **(Figure 4D, left)**. Additionally, fibrosis-related genes including *COL1A1* and *COL2A1* were significantly inhibited by SAHA **(Figure 4D, right)**. Together, these results suggested that SAHA improves hepatic injury in our hepatocyte-specific *Asah1*-deficient PD model, especially inflammation and fibrosis.

To evaluate the impairment of LSEC and the therapeutic effect, hepatic genes related to LSEC phenotype (LSEC differentiation, capillarization, and vascular tone dysregulation) were measured by qPCR **(Figure 4E)**. Results showed that hepatocyte-specific *Asah1*-deficiency with PD did not significantly change hepatic gene expression levels of LSEC differentiation markers (*Lymphatic vessel endothelial hyaluronan receptor 1 [LYVE1]* and *Kinase insert domain receptor* [*KDR]*), while their expression were not significantly altered by SAHA **(Figure 4E, left)**. Hepatocyte- specific *Asah1*-deficiency with PD up-regulated *CD34* (LSEC capillarization marker) and *endothelin-1* (*EDN1*, vasoconstrictor) expression. Meanwhile, *plasmalemma vesicle-associated protein* (*PLVAP)* and *von Willebrand factor (VWF)* (both capillarization markers), as well as *nitric oxide synthase 3* (*NOS3,* mediates eNOS/NO production) remained unchanged. Notably, SAHA significantly inhibited *CD34* and *EDN1* expression, and decreased the basal level of *VWF* (**Figure 4E, middle and right)**. Protein expression levels of ICAM-1 (endothelial inflammation marker), CD34, and ET-1 in LSECs were evaluated by colocalization with IB4 (endothelial marker) **(Figure 4F)**. Consistently, SAHA reduced the colocalized area of ICAM-1, CD34, and ET-1, indicating improved LSEC inflammation, capillarization, and vasoconstriction.

Portal venous hemodynamics and cardiac function were assessed by ultrasonography at weeks 12 and 20. At week 20, SAHA significantly improved portal venous velocities, including V_max_, V_min_, and V_mean_, while PI remained unchanged **(Figure 4G-H)**. Meanwhile, hepatocyte-specific *Asah1*-deficient mice with PD developed systolic cardiac dysfunction, manifested by impaired ventricular contractility (decreased EF and stroke volume), incomplete ventricular emptying (increased LVID;s and LV Vol;s), and preserved diastolic filling capacity (unchanged LVID;d and LV Vol;d) **(Figure 4I-J)**. However, at the current dose (15 mg/kg/day), SAHA did not improve cardiac function despite hepatic and portal hemodynamic benefit, suggesting that its effect at this dose is predominantly liver- and LSEC-associated. In addition, portal hemodynamics and cardiac function were both preserved at week 12 **(Supplementary Figure 1)**, indicating the occurrence of abnormalities between weeks 12 and 20, coinciding with disease progression.

### *Asah1*-deficient hepatocytes induce LSEC dysfunction through paracrine HMGB1 signaling

The aggravation of LSEC dysfunction observed in hepatocyte-specific *Asah1*-deficient mice indicated a potential intercellular crosstalk between *Asah1*-deficient hepatocytes and LSECs. To identify the potential hepatocyte-derived paracrine mediator responsible for LSEC dysfunction, we first reviewed the established hepatocyte-derived factors implicated in intercellular crosstalk with other hepatic cell types (HSCs, LSECs, immune cells) during fibrotic liver disease **(Figure 5A)**[52–70] .

**Figure 5.**
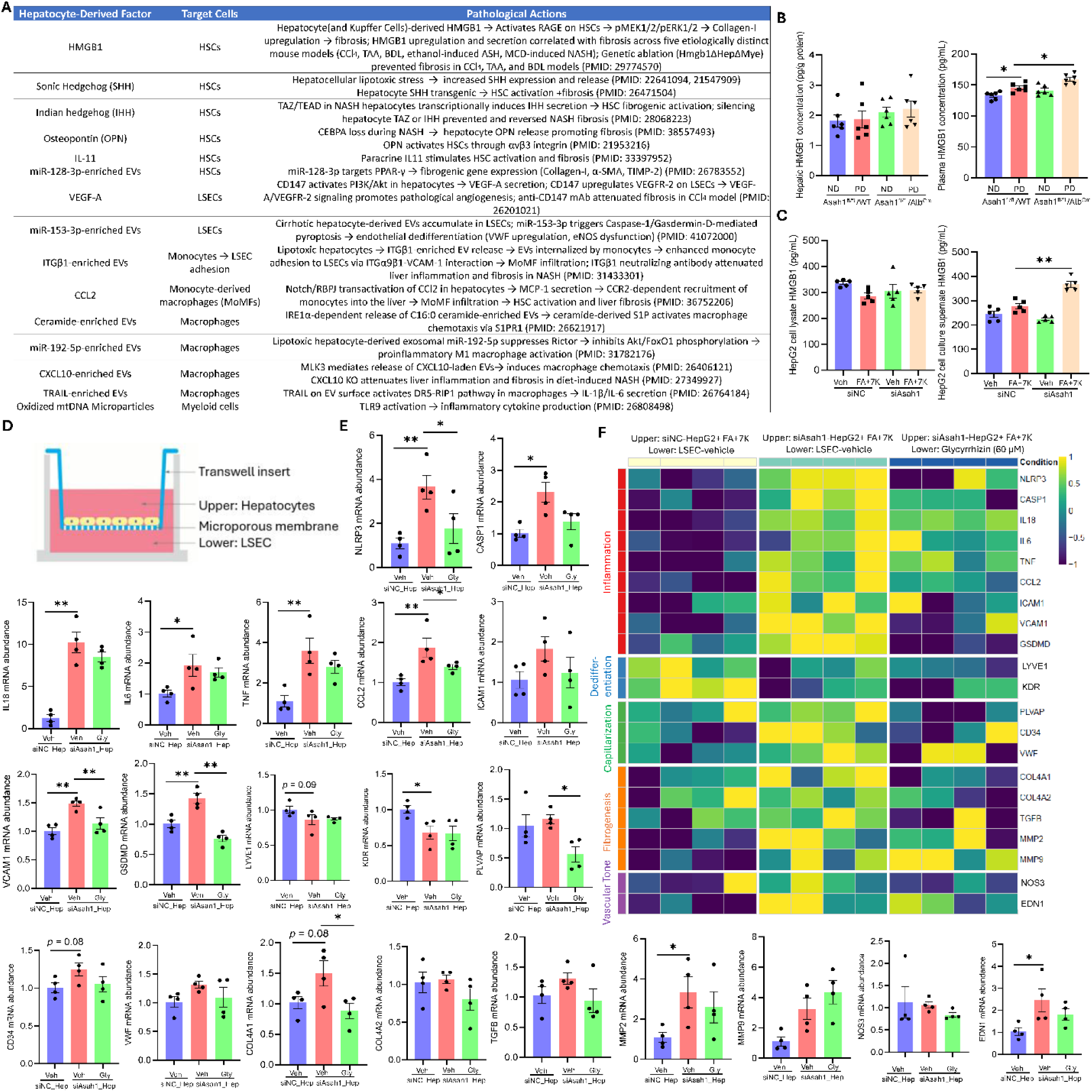
*Asah1*-deficient hepatocytes induce LSEC injury through paracrine HMGB1 signaling. **(A)** Summary of hepatocyte-derived factors in intercellular crosstalk during NAFLD pathogenesis. **(B)** *Asah1* floxed (*Asah1*^fl/fl^/WT) mice and *Asah1*^fl/fl^/*Alb*^Cre^ mice (hepatocyte-specific deletion of *Asah1*) were fed with PD for 20 weeks. Hepatic and plasma HMGB1 levels were measured by ELISA. Hepatic HMGB1 level was normalized to total protein. **(C)** HepG2-hepatocytes were transfected with siNC or si*ASAH1* for 48 hours, followed by treatment with vehicle or a lipid mixture consisting of free FAs (300 μM; OA: PA = 2:1) and 7K (40 μM) for 24 hours. Cell lysate and supernatant HMGB1 levels were quantified by ELISA. **(D)** Transwell co-culture schematic. HepG2 hepatocytes were cultured in the upper chamber, transfected with siNC or *siASAH1* for 48 hours. Primary LSECs were cultured separately in the lower chamber. After transfection, co-culture was initiated with FA + 7K treatment in the upper chamber and vehicle or glycyrrhizin (60 μM, HMGB1 inhibitor) in the lower chamber for 24 hours. **(E)** qPCR quantification (normalized to *HPRT1*) and **(F)** heatmap of LSEC marker expression. Mean ± SEM; n = 4. \**p* < 0.05, \*\**p* < 0.01. Abbreviations: HMGB1, high mobility group box 1; HSCs, hepatic stellate cells; RAGE, receptor for advanced glycation end products; pMEK1/2, phosphorylated mitogen-activated protein kinase kinase 1/2; pERK1/2, phosphorylated extracellular signal-regulated kinase 1/2; CCl₄, carbon tetrachloride; TAA, thioacetamide; BDL, bile duct ligation; ASH, alcoholic steatohepatitis; MCD, methionine-choline-deficient; NASH, nonalcoholic steatohepatitis; Hmgb1ΔHepΔMye, hepatocyte and myeloid cell-specific HMGB1 knockout; SHH, Sonic hedgehog; IHH, Indian hedgehog; TAZ, transcriptional co-activator with PDZ-binding motif (WWTR1); TEAD, TEA domain transcription factor; OPN, osteopontin; CEBPA, CCAAT/enhancer-binding protein alpha; EVs, extracellular vesicles; PPAR-γ, peroxisome proliferator-activated receptor gamma; α-SMA, alpha-smooth muscle actin; TIMP-2, tissue inhibitor of metalloproteinase 2; VEGF-A, vascular endothelial growth factor A; PI3K, phosphoinositide 3-kinase; Akt, protein kinase B; VEGFR-2, vascular endothelial growth factor receptor 2; mAb, monoclonal antibody; VWF, von Willebrand factor; eNOS, endothelial nitric oxide synthase; ITGβ1, integrin beta 1; ITGα9β1, integrin alpha 9 beta 1; VCAM-1, vascular cell adhesion molecule 1; MoMF, monocyte-derived macrophage; CCL2, C-C motif chemokine ligand 2; RBPJ, recombination signal binding protein for immunoglobulin kappa J region; MCP-1, monocyte chemoattractant protein-1; CCR2, C-C motif chemokine receptor 2; IRE1α, inositol-requiring enzyme 1 alpha; S1P, sphingosine-1-phosphate; S1PR1, sphingosine-1-phosphate receptor 1; FOXO1, forkhead box O1; CXCL10, C-X-C motif chemokine ligand 10; MLK3, mixed lineage kinase 3; TRAIL, TNF-related apoptosis-inducing ligand; DR5, death receptor 5; RIP1, receptor-interacting protein 1; mtDNA, mitochondrial DNA; TLR9, Toll-like receptor 9; FA, fatty acid; 7K, 7-ketocholesterol; siNC, negative control siRNA; siASAH1, ASAH1 siRNA; Gly, Glycyrrhizin; HPRT1, hypoxanthine phosphoribosyltransferase; NLRP3, NLR family pyrin domain containing 3; CASP1, caspase-1; ICAM1, intercellular adhesion molecule 1; GSDMD, gasdermin D; LYVE1, lymphatic vessel endothelial hyaluronan receptor 1; KDR, kinase insert domain receptor; PLVAP, plasmalemma vesicle-associated protein; COL4A1, collagen type IV alpha 1 chain; COL4A2, collagen type IV alpha 2 chain; TGF-β, transforming growth factor beta; MMP2, matrix metallopeptidase 2; MMP9, matrix metallopeptidase 9; NOS3, nitric oxide synthase 3; ET-1, endothelin-1.

Among these factors, HMGB1 was of particular interest as it is a well-characterized damage-associated molecular pattern (DAMP) and has been demonstrated to be released from stressed hepatocytes in five etiologically distinct mouse liver fibrosis models (including hepatotoxins [CCl_4_ or thioacetamide, TAA]-induced liver fibrosis, bile duct ligation-induced cholestatic liver fibrosis, ethanol-induced alcoholic steatohepatitis, and methionine-choline-deficient [MCD] diet-induced NASH) [52]. To determine whether HMGB1 is elevated in the *Asah1*-deficient model, we measured hepatic and plasma HMGB1 levels in *Asah1*^fl/fl^/WT and *Asah1*^fl/fl^/*Alb*^Cre^ mice fed ND or PD for 20 weeks **(Figure 5B)**. Plasma HMGB1 was significantly elevated in PD mice and further increased by *Asah1* deficiency, while hepatic HMGB1 level remained unchanged. Similarly, in cell supernatant of cultured HepG2 hepatocytes, HMGB1 level was modestly elevated by FA + 7K treatment in WT cells and significantly induced by *ASAH1* siRNA transfection **(Figure 5C)**. Meanwhile, HMGB1 level in cell lysate remained unchanged. These results indicated that *Asah1* deficiency promotes HMGB1 release, rather than intrahepatocellular accumulation.

To test whether hepatocyte-derived HMGB1 drives LSEC dysfunction, we established a Transwell co-culture system with si*ASAH1*- or siNC-transfected HepG2 hepatocytes (both treated with FA+7K) in the upper chamber and primary LSECs (treated with vehicle or Gly, a known HMGB1 antagonist) in the lower chamber **(Figure 5D)**. After FA + 7K treatment, LSEC gene expression was assessed by qPCR across five functional domains **(Figure 5E-F)**. Co-culture with si*ASAH1*-hepatocytes under lipotoxic stress induced broad LSEC dysfunction across multiple functional domains. In the inflammation domain, si*ASAH1*-hepatocytes significantly upregulated pro-inflammatory mediators (*IL18, IL6, TNF, CCL2*), NLRP3 inflammasome and pyroptosis components (*NLRP3, CASP1, GSDMD*), and the adhesion molecule *VCAM1*. On the other hand, *ICAM1* showed an increasing trend without reaching significance. LSEC dedifferentiation was evidenced by significantly decreased *KDR* expression, with *LYVE1* showing a decreasing trend. Capillarization markers *CD34* and *VWF* showed increasing trends, while PLVAP remained unchanged. In the fibrogenesis domain, *MMP2* was significantly upregulated, with *COL4A1*, *TGFB1*, and *MMP9* showing increasing trends; *COL4A2* remained unchanged. For vascular tone, *EDN1* was significantly increased, while *NOS3* was unaffected.

Importantly, treatment of LSECs with Gly (60 μM), a well-established HMGB1 inhibitor, partially reversed these changes. Gly significantly attenuated *NLRP3, GSDMD, VCAM1*, and *COL4A1* expression, with modest reductions in several other inflammatory and capillarization markers **(Figure 5E-F).** However, Gly did not reverse *IL6, LYVE1, KDR, MMP9*, or *NOS3* expression. Together, these results indicated that HMGB1 is a major but not exclusive mediator of hepatocyte-to-LSEC paracrine injury, and that additional hepatocyte-derived factors likely contribute to the full spectrum of LSEC dysfunction.

### SAHA reverses HMGB1-induced LSEC dysfunction in vitro

Having established HMGB1 as a key paracrine mediator of LSEC injury, we next tested whether SAHA directly protects LSECs against HMGB1-induced dysfunction **(Figure 6A-B)**. HMGB1 treatment induced comprehensive LSEC dysfunction, recapitulating the pattern observed in the Transwell co-culture system. In the inflammation domain, HMGB1 significantly upregulated *CASP1* and *CCL2,* with *IL18, IL6, TNF, NLRP3, GSDMD,* and *ICAM1* showing increasing trends. HMGB1 also promoted LSEC dedifferentiation (*KDR* showing decreasing trend), capillarization (*PLVAP* showing increasing trend), fibrogenesis (increased *TGFB1*, with *COL4A1, COL4A2, MMP9* showing trends), and vascular tone dysregulation (decreased *NOS3* trend). Importantly, SAHA dose-dependently improved most of these HMGB1-induced changes. At the medium (5 μM) or high dose (10 μM), SAHA significantly attenuated *CASP1, IL18, IL6, TNF, CCL2, ICAM1, VCAM1, GSDMD, CD34, COL4A1, COL4A2, MMP2, TGFB1,* and *EDN1* gene expression. *NLRP3, VWF* and *MMP9* showed improving trends. No effects on *PLVAP, LAMA1*, and *NOS3* were observed in SAHA-treated groups. It should be noted that the basal levels of dedifferentiation (*LYVE1* and *KDR*) were both down-regulated by SAHA, raising a concern that SAHA may independently promote LSEC dedifferentiation.

**Figure 6.**
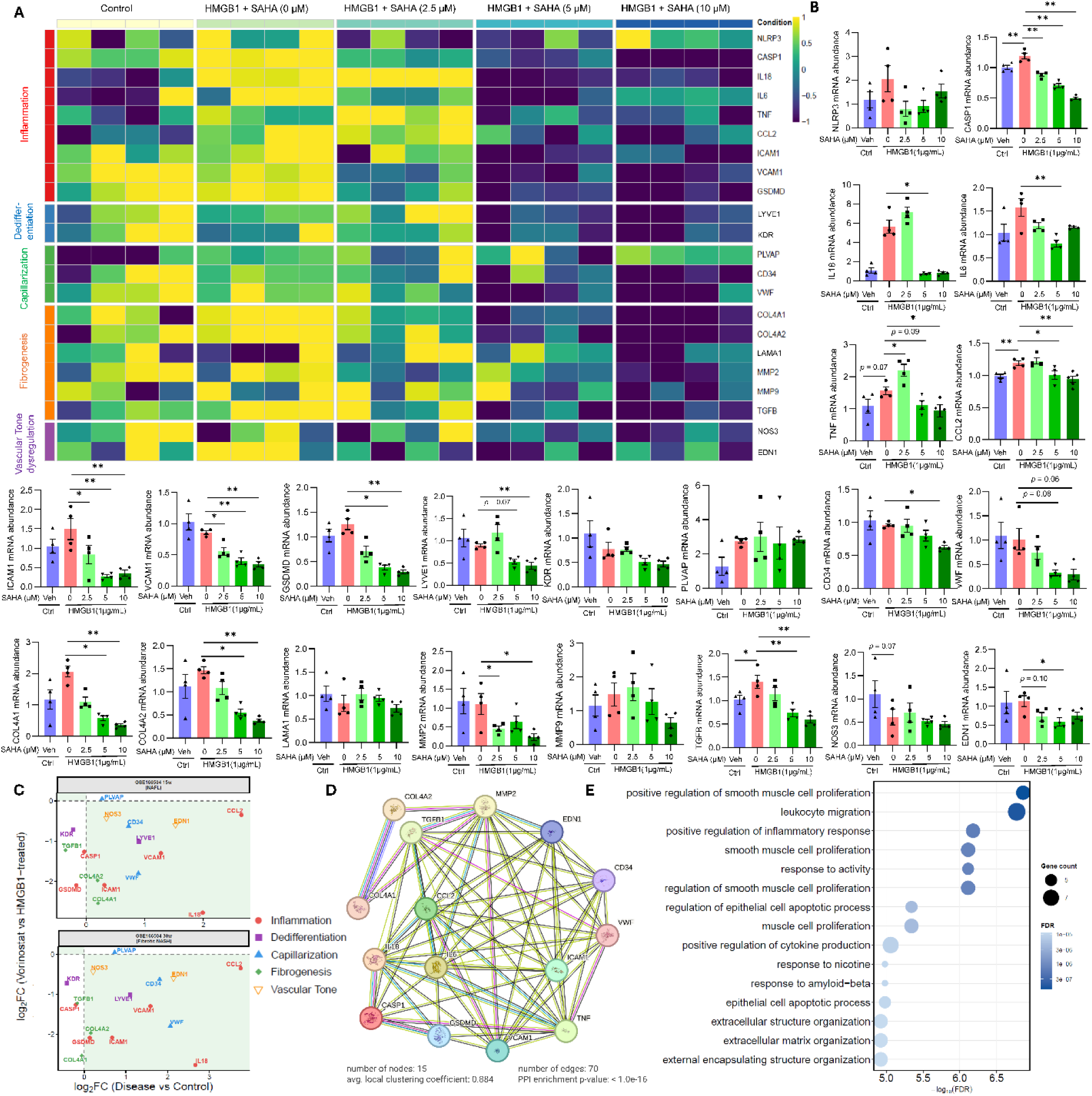
SAHA attenuates HMGB1-induced primary LSEC dysfunction in vitro. (A-B) Heatmap and qPCR of primary LSECs treated with vehicle or HMGB1 (1 μg/mL) ± SAHA (0, 2.5, 5, 10 μM) across five domains: Inflammation, Dedifferentiation, Capillarization, Fibrogenesis, and Vascular tone dysregulation. Results were normalized to *HPRT1*. **(C)** Correlation plot comparing disease-induced LSEC gene changes (x-axis, disease vs. control log₂FC) with SAHA reversal effects (y-axis, SAHA-treated vs. HMGB1-treated log₂FC) in simple steatosis NAFLD (HFHFD 15 weeks) and fibrotic NASH (HFHFD 30 weeks) from GSE166504. **(D)** PPI network and **(E)** GO enrichment analysis of 15 genes reversed by SAHA. Mean ± SEM; n = 4. \**p* < 0.05, \*\**p* < 0.01. Abbreviations: NLRP3, NLR family pyrin domain containing 3; CASP1, caspase-1; IL18, interleukin-18; IL6, interleukin-6; TNF, tumor necrosis factor; CCL2, C-C motif chemokine ligand 2; ICAM1, intercellular adhesion molecule 1; VCAM1, vascular cell adhesion molecule 1; GSDMD, gasdermin D; LYVE1, lymphatic vessel endothelial hyaluronan receptor 1; KDR, kinase insert domain receptor; PLVAP, plasmalemma vesicle-associated protein; CD34, cluster of differentiation 34; VWF, von Willebrand factor; COL4A1, collagen type IV alpha 1 chain; COL4A2, collagen type IV alpha 2 chain; LAMA1, laminin subunit alpha 1; MMP2, matrix metallopeptidase 2; MMP9, matrix metallopeptidase 9; TGFB1, transforming growth factor beta1; NOS3, nitric oxide synthase 3; EDN1, endothelin-1; FC, fold change; PPI, protein-protein interaction; GO, Gene Ontology; FDR, false discovery rate.

To assess whether the in vitro reversal pattern corresponds to disease-relevant gene changes, we correlated the SAHA treatment effect (log₂ fold change [log₂FC], SAHA-treated vs. HMGB1-treated) with the disease-induced LSEC gene changes (log₂FC, disease vs. control) from GSE166504 **(Figure 6C).** Genes reversed by SAHA (10 μM) with clinical relevance in both simple steatosis (HFHFD 15 weeks) and fibrotic NASH (HFHFD 30 weeks) include *IL18, CCL2, VCAM1, ICAM1, CD34, VWF, COL4A1, COL4A2,* and *EDN1*. The correlation plot demonstrated that disease-upregulated genes were predominantly reversed by SAHA (falling into the lower-left diagonal), with similar reversal magnitudes in simple steatosis and fibrotic NASH stages.

Notably, SAHA-mediated *LYVE1* and *KDR* suppression may limit its endothelial-protective scope. PPI network analysis of the 15 SAHA-reversed genes revealed a highly interconnected network (70 edges, PPI enrichment p < 1.0e-16) **(Figure 6D)**, and GO enrichment analysis further confirmed that SAHA-mediated processes include inflammatory response regulation, leukocyte migration, cytokine production, and extracellular matrix organization, covering both inflammation and fibrosis pathways **(Figure 6E).**

### SAHA does not antagonize HMGB1-receptor signaling in the hepatocyte-LSEC axis

Given that SAHA reversed HMGB1-induced LSEC dysfunction, we next investigated whether this protection involves direct antagonism of HMGB1-receptor interactions. We first characterized the HMGB1-LSEC receptor signaling landscape across disease stages using scRNA-seq data. Interaction strength was quantified for nine known HMGB1 receptors across mouse (GSE166504: chow, HFHFD 15 weeks, HFHFD 30 weeks) and human (GSE136103: healthy, cirrhotic) datasets **(Figure 7A).** Among these receptors, TLR4 and THBD showed high expression across both mouse and human LSECs, while CXCR4 was prominently expressed in human LSECs. Interestingly, AGER (or RAGE), one of the classic HMGB1 receptors, is lowly expressed in both mouse and human LSECs, suggesting that HMGB1 signaling in LSECs is primarily mediated through TLR4/CXCR4/THBD, rather than the classical RAGE pathway. Chord diagrams further illustrated the shifting receptor contributions to total HMGB1 signaling at each disease stage **(Figure 7B)**.

**Figure 7.**
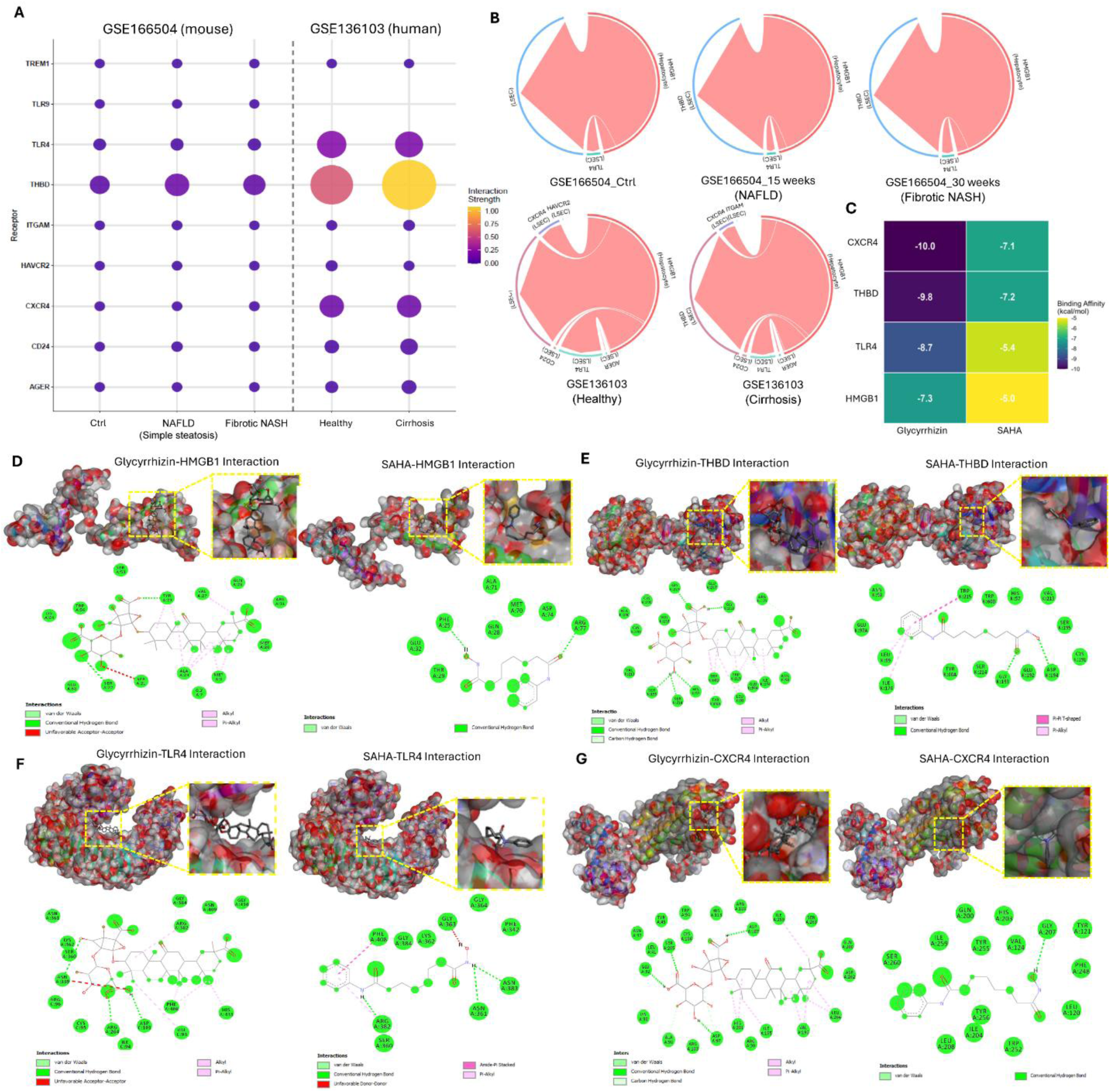
SAHA does not antagonize HMGB1-receptor signaling in the hepatocyte-LSEC axis. **(A)** Dot plot of hepatocyte HMGB1 to known LSEC receptor interaction strength across disease stages in mouse (GSE166504: Chow, NAFLD simple steatosis 15 weeks, fibrotic NASH 30 weeks) and human (GSE136103: Healthy, Cirrhotic) scRNA-seq data. Dot size and color encode interaction strength. **(B)** Chord diagrams showing receptor-level contribution to total HMGB1 signaling at each disease stage. Chord width is proportional to interaction strength. **(C)** Binding affinity (kcal/mol) heatmap for Gly and SAHA against HMGB1, CXCR4, THBD, and TLR4. More negative values indicate stronger binding. **(D-G)** 3D docking poses (top) and 2D interaction diagrams (bottom) for Gly (left) and SAHA (right) with (D) HMGB1 A+B box (PDB: 2YRQ), (E) thrombin/THBD complex (PDB: 1HLT), (F) TLR4/MD-2 complex (PDB: 3FXI), and (G) CXCR4 7-TM bundle core (PDB: 3ODU). Yellow boxes highlight binding pockets. Interaction types are color-coded in the 2D diagrams. Abbreviations: Gly, glycyrrhizin; TREM1, triggering receptor expressed on myeloid cells 1; TLR9, Toll-like receptor 9; TLR4, Toll-like receptor 4; THBD, thrombomodulin; ITGAM, integrin subunit alpha M; HAVCR2, hepatitis A virus cellular receptor 2; CXCR4, C-X-C chemokine receptor type 4; CD24, cluster of differentiation 24; AGER (or RAGE), advanced glycosylation end-product specific receptor; MD-2, myeloid differentiation factor 2; PDB, Protein Data Bank; 7-TM, seven-transmembrane.

To determine whether SAHA directly competes with HMGB1 at its receptor binding sites, we performed molecular docking of both SAHA and glycyrrhizin (positive control) against HMGB1 and three key LSEC receptors: TLR4, THBD, and CXCR4 **(Figure 7C-G)**. Glycyrrhizin exhibited consistently stronger binding affinities than SAHA across all four targets (CXCR4: -10.0 vs -7.1; THBD: -9.8 vs -7.2; TLR4: -8.7 vs -5.4; HMGB1: -7.3 vs -5.0 kcal/mol) (**Figure 7C**), likely attributable to its larger molecular size enabling more extensive surface contact with binding pockets (**Figure 7D-G**). Detailed interaction profiles are provided in **Supplementary Tables 2-3**. These results suggest that SAHA is unlikely to function as a direct HMGB1-receptor antagonist (especially due to its poor binding affinities with HMGB1 and TLR4), and that its LSEC-protective effect is more likely mediated through intracellular mechanisms.

### SAHA modulates endothelial transcription factor profiles and upregulates KLF2 in primary LSECs

Since direct HMGB1-receptor antagonism has been excluded, we next investigated the intracellular mechanism underlying SAHA’s LSEC-protective effect. Considering that SAHA is a pan-HDAC inhibitor that broadly modulates gene transcription through histone acetylation, we hypothesized that its protective effect may involve reprogramming of the endothelial transcription factor (TF) landscape. To test this, we analyzed publicly available RNA-seq data from SAHA-treated HAECs (GSE54912).

GSVA across 12 Hallmark pathways in SAHA-treated HAECs (**Figure 8A**) showed that TGF-β signaling and ROS pathway were significantly suppressed by SAHA, while Inflammatory response and IL6/JAK-STAT3 pathways showed decreasing trends. Differentially expressed TF analysis identified widespread transcriptomic changes in SAHA-treated HAECs **(Figure 8B)**. Among these, SAHA significantly upregulated endothelial-protective TFs including *KLF2, KLF4, NR4A1, NR4A2, PPARA* and *PPARD,* while simultaneously downregulating pro-fibrotic and pro-inflammatory TFs including *NFKB1, SMAD3*, and the mechanotransduction effectors *YAP1* and *WWTR1 (TAZ)* **(Figure 8C)**.

**Figure 8.**
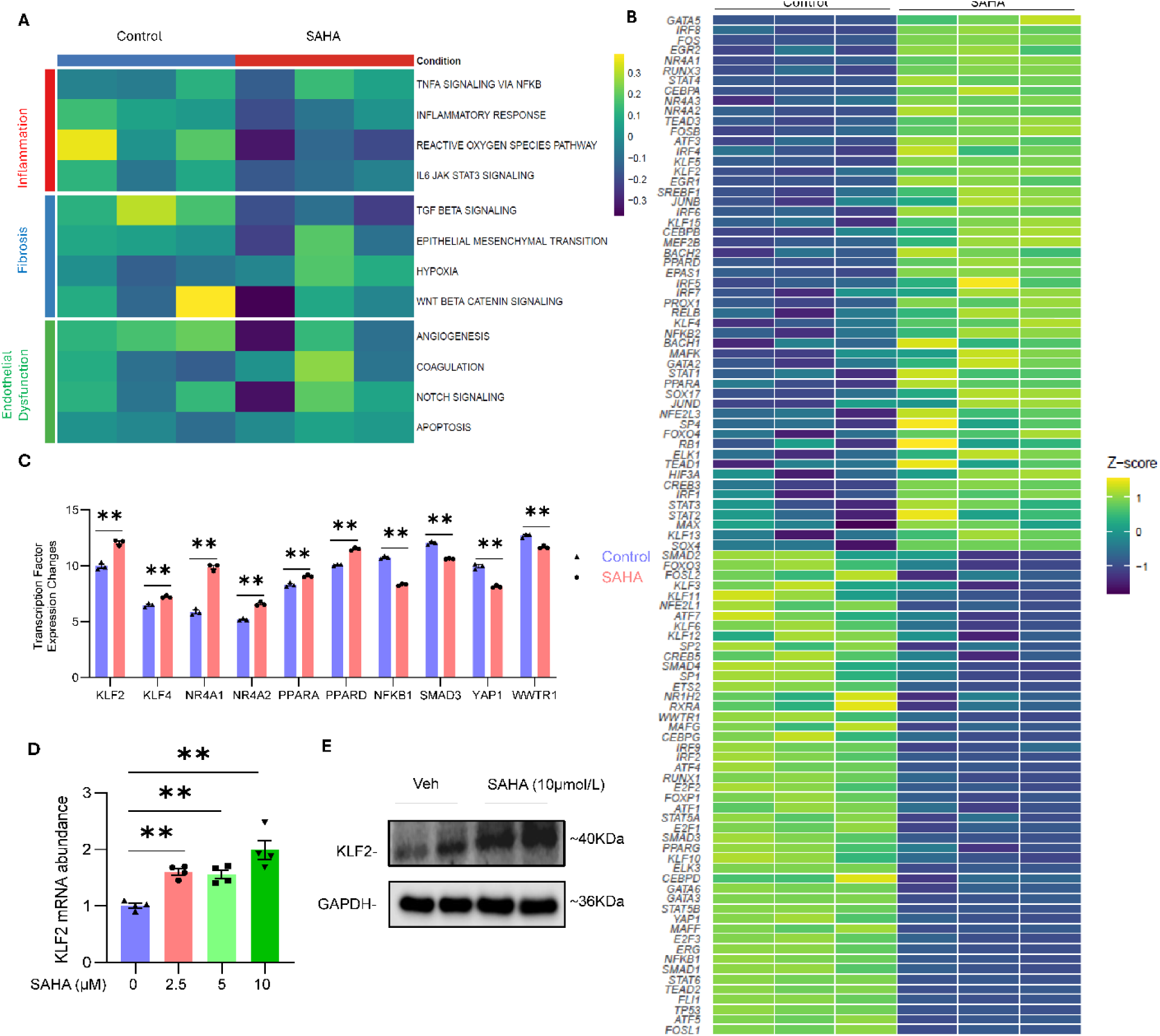
SAHA modulates a broad panel of transcription factors in endothelial cells and upregulates KLF2 in primary LSECs. **(A)** GSVA heatmap of 12 Hallmark pathways across 3 functional categories in SAHA-treated HAEC (bulk RNA-seq, GSE54912). **(B)** Differentially expressed TF heatmap in control vs SAHA-treated HAEC. **(C)** Expression fold change of protective and detrimental TFs in SAHA-treated HAEC. **(D)** *KLF2* mRNA expression in primary LSECs treated with SAHA (0, 2.5, 5, 10 μM), normalized to HPRT1. **(E)** Representative Western blot of KLF2 protein in primary LSECs treated with vehicle or SAHA (10 μM), normalized to GAPDH. Mean ± SEM, n = 3/group (RNA-seq), n = 4 (LSEC validation). \*\**p* < 0.01. Abbreviations: HAEC, human aortic endothelial cell; TF, transcription factor; KLF2, Krüppel-like factor 2; KLF4, Krüppel-like factor 4; NR4A1, nuclear receptor subfamily 4 group A member 1; NR4A2, nuclear receptor subfamily 4 group A member 2; PPARA, peroxisome proliferator-activated receptor alpha; PPARD, peroxisome proliferator-activated receptor delta; NFKB1, nuclear factor kappa B subunit 1; SMAD3, SMAD family member 3; YAP1, Yes-associated protein 1; WWTR1, WW domain-containing transcription regulator 1; HPRT1, hypoxanthine phosphoribosyltransferase; GAPDH, glyceraldehyde-3-phosphate dehydrogenase.

Among these TFs, we chose KLF2 to validate in primary LSECs due to its well-established role as a TF that mediates LSEC protection as well as its consistent downregulation in our LSEC gene signature analysis **(Figure 2G)**. To validate *KLF2* upregulation in LSECs, primary mouse LSECs were treated with SAHA (0, 2.5, 5, or 10 μM) for 24 hours. SAHA dose-dependently increased *KLF2* mRNA expression, reaching significance at all the three doses **(Figure 8D).** Western blot confirmed increased KLF2 protein expression at 10 μM **(Figure 8E).**

### SAHA improves LSEC dysfunction in CCl_4_-induced liver fibrosis and TNF-α-induced inflammation models

To test whether the LSEC-protective effect of SAHA extends beyond the *Asah1*-deficient Paigen diet model, we employed CCl_4_-induced liver fibrosis as an etiologically distinct injury model in WT C57BL/6J mice **(Figure 9A)**. SAHA treatment did not affect body weight but significantly reduced circulating ALT and AST levels in CCl_4_-treated mice **(Figure 9B).** Histological and immunofluorescence analysis confirmed that SAHA attenuated CCl_4_-induced hepatic injury, including reduced collagen deposition (Sirius Red, Collagen I), decreased apoptosis (TUNEL), and diminished immune cell infiltration (CD45, CD11b) **(Figure 9C).** Co-immunofluorescence staining with IB4 demonstrated that SAHA increased sinusoidal KLF2 expression, reduced ICAM-1 and VCAM-1, decreased CD34, and attenuated ET-1 expression in LSECs **(Figure 9C).** Portal venous hemodynamics were then evaluated by ultrasonography **(Figure 9D-E).** CCl_4_ treatment significantly decreased V_max_ and showed decreasing trend in V_min_ and V_mean_. Interestingly, unlike the PD model, CCl_4_ treatment significantly decreased PI, suggesting distinct hemodynamic responses between the two models. Nonetheless, SAHA did not significantly improve portal venous hemodynamics in CCl_4_ model.

**Figure 9.**
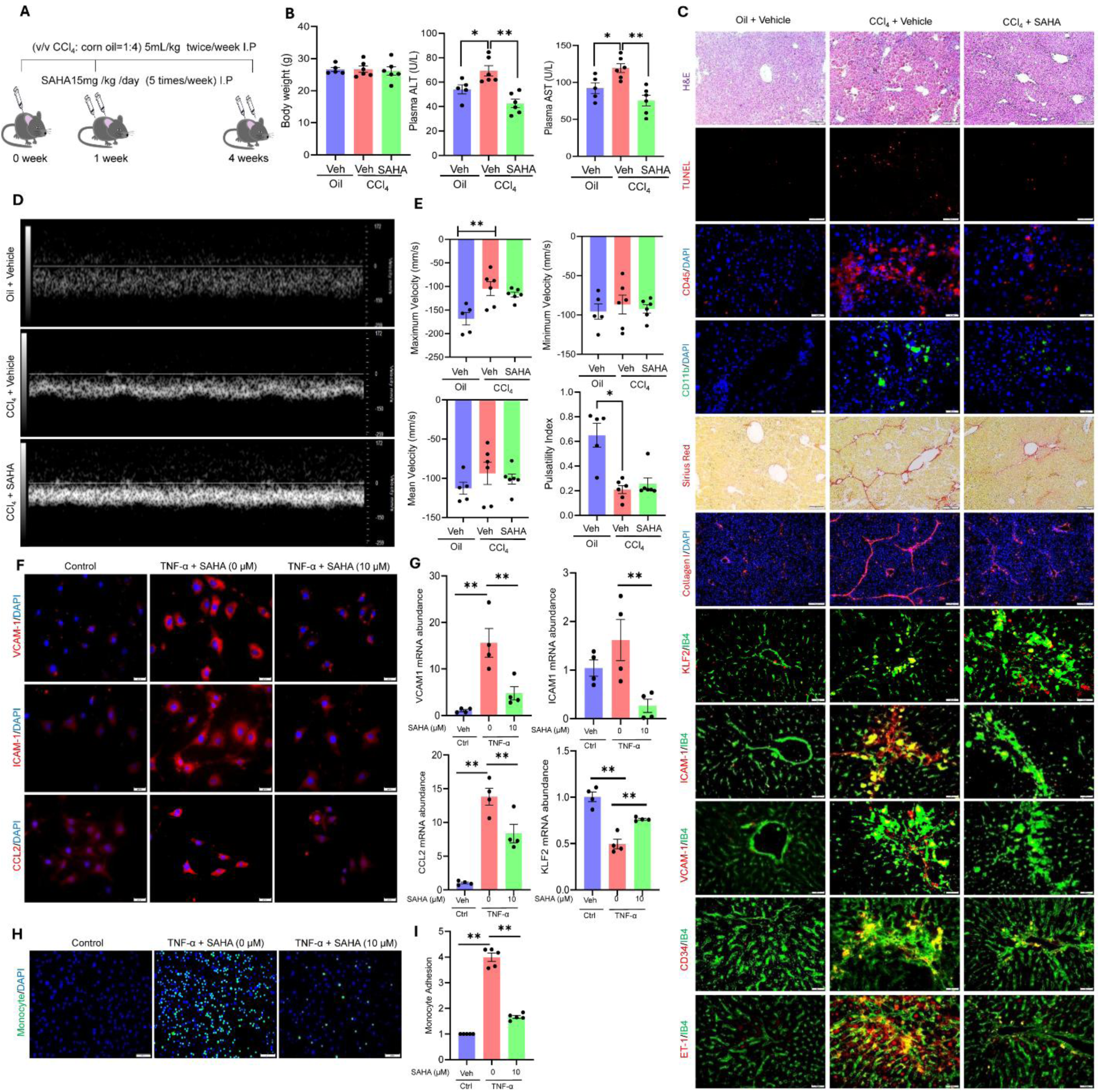
SAHA improves LSEC dysfunction in CCl₄-induced liver fibrosis and TNF-α-induced inflammation models. **(A)** CCl_4_ model design. WT C57BL6/J mice were administered with vehicle or CCl₄ (v/v CCl₄: corn oil = 1:4, 5 mL/kg, i.p., twice weekly) from week 0, with SAHA treatment (15 mg/kg/day, i.p., 5 times/week) initiated at week 1, and sacrificed at week 4. Portal venous hemodynamics was assessed by ultrasonography at week 4. **(B)** Mouse body weight (g), plasma ALT (U/L), and plasma AST (U/L). **(C)** Paraffin liver sections (4-μm-thick) were used for pathological staining (H&E, Sirius Red). Frozen liver sections (8-μm-thick) were used for immunofluorescence staining (TUNEL, CD45/DAPI, CD11b/DAPI, Collagen I/DAPI, KLF2/IB4, ICAM-1/IB4, VCAM-1/IB4, CD34/IB4, ET-1/IB4). **(D-E)** Representative portal vein Doppler waveforms (week 4) and hemodynamic quantification including maximum velocity (mm/s), minimum velocity (mm/s), mean velocity (mm/s), and pulsatility index. **(F-G)** Primary LSECs were treated with TNF-α (40 ng/mL) ± SAHA (0 or 10 μM), followed by immunofluorescence staining (VCAM1, ICAM1, or CCL2 with DAPI) and qPCR quantification of *VCAM1, ICAM1, CCL2*, and *KLF2* (normalized to *HPRT1*). **(H-I)** Representative fluorescence images and quantification of monocyte adhesion to TNF-α-stimulated LSECs ± SAHA (10 μM). Mean ± SEM; n = 5-6/group (in vivo), n = 4 (in vitro). \**p* < 0.05, \*\**p* < 0.01. Abbreviations: TNF-α, tumor necrosis factor alpha; i.p., intraperitoneal; ALT, alanine aminotransferase; AST, aspartate aminotransferase; H&E, hematoxylin and eosin; TUNEL, terminal deoxynucleotidyl transferase-mediated dUTP nick-end labeling; DAPI, 4’,6-diamidino-2-phenylindole; IB4, isolectin B4; KLF2, Krüppel-like factor 2; ICAM-1, intercellular adhesion molecule 1; VCAM-1, vascular cell adhesion molecule 1; ET-1, endothelin-1; CCL2, C-C motif chemokine ligand 2; HPRT1, hypoxanthine phosphoribosyltransferase.

To further dissect the direct endothelial-protective effect of SAHA independent of hepatocyte-derived paracrine signaling, we treated primary LSECs with TNF-α (40 ng/mL) with or without SAHA (10 μM) for 12 hours **(Figure 9F-G)**. TNF-α robustly upregulated the protein expression of VCAM-1, ICAM-1, and CCL2, as shown by IF staining **(Figure 9F).** qPCR analysis **(Figure 9G)** showed that TNF-α significantly increased *VCAM1* and *CCL2* gene expression, while *ICAM1* showed an increasing trend. Meanwhile, TNF-α significantly decreased *KLF2* gene expression. SAHA significantly reversed all four markers: attenuating *VCAM1, ICAM1,* and *CCL2* induction while restoring *KLF2* expression **(Figure 9F-G).** Furthermore, monocyte adhesion assay was performed as a functional endpoint. TNF-α-stimulated LSECs showed markedly increased J774 macrophage adhesion, which was significantly reduced by SAHA co-treatment **(Figure 9H-I)**, providing functional evidence that SAHA attenuates LSEC activation and leukocyte recruitment.

## DISCUSSION

In this study, we established a systematic LSEC-focused computational drug screening platform for fibrotic liver diseases. First, by focusing specifically on LSEC transcriptomic profiles rather than whole-liver or hepatocyte-specific data, the workflow prioritized cell-type relevance. Next, the tiered gene signature selection strategy was designed to balance statistical stability with biological interpretability by combining stability selection, LLM-assisted literature curation, and human expert review. Moreover, the cross-stage (simple steatosis NAFLD, fibrotic NASH, cirrhosis) and cross-species (mouse and human) integration enables identification of candidates with broad-spectrum LSEC-protective potential. Since LSEC-specific perturbational profiles are not available in CMap, HUVEC profiles were used as the closest endothelial perturbation resource. This substitution may not fully recapitulate LSEC-specialized features such as fenestration. Accordingly, HUVEC-based CMap results were not interpreted as direct evidence of LSEC efficacy, but as a prioritization step for downstream validation. The independent support for several candidates, together with experimental validation of SAHA in primary LSECs and two fibrosis models, supports the utility of this pragmatic screening strategy.

Another advantage of our platform is the integration of gene safety assessment (DepMap, COSMIC, ClinVar, literature review) before CMap screening, which minimizes the dilution of risky genes (e.g., developmentally essential or oncogenic genes). This is particularly important due to the inconsistencies revealed by GSVA across disease stages. Specifically, while mouse NAFLD and fibrotic NASH datasets (GSE166504, GSE140994) showed similar patterns, the human cirrhosis dataset (GSE136103) exhibited a distinct pattern with negative enrichment of inflammation pathways. This divergent pattern likely reflects stage- and species- dependent remodeling and may be attributed to a phenomenon named “burnt-out” steatohepatitis in end-stage cirrhosis, when the early inflammatory and steatotic features diminish as fibrosis advances, leaving a fibrotic liver with attenuated inflammatory markers [71]. Our gene safety assessment mitigated the risk of reversing potentially adaptive gene changes at cirrhosis stage and ensured that only safely targetable genes were included for CMap query.

However, several limitations of our computational platform should be mentioned. First, LLM outputs may lack guaranteed reproducibility, as responses may vary across sessions, model versions, or prompt formulations. To mitigate this, the LLM in our platform was restricted to a Tier 2 advisory role. Specifically, LLM was used to assist retrieval-augmented literature review, with all recommendations subject to mandatory human expert validation. Second, the limited biological replicates in bulk and scRNA-seq data may introduce potential overfitting issues. Therefore, in this study, we chose stability selection for its robustness in high-dimensional, low-sample-size settings [27, 72, 73], and applied it for feature selection instead of predictive modeling, with input restricted to pre-filtered DEGs (FDR < 0.25). The ultimate validation of platform output relied on independent experimental and clinical evidence rather than computational prediction accuracy.

Using the computational platform, we identified 6 clinical-stage and 8 preclinical drug candidates with LSEC-protective potential, and chose SAHA as our drug of interest. In line with our research, Guan et al. recently reported that an AI co-scientist system also identified SAHA (HDAC inhibitor) and several BET inhibitors (XMD8-85, XMD8-92, and I-BET151 by our platform; ZEN-3694, I-BET762, and JQ-1 by Guan et al.)[48]. The difference is that their methods employed AI-assisted hypothesis generation that synthesized existing literature to directly recommend drug candidates, whereas our platform used LLM solely for gene signature curation, with drug candidate identification driven by unbiased CMap transcriptomic screening. The convergent identification of HDAC and BET inhibitors by two methodologically distinct AI-assisted workflows provides independent support for the biological plausibility of these candidate classes.

Wang et al. previously reported that SAHA alleviates CCl_4_-induced liver fibrosis in rats by restoring histone acetylation, upregulating the inhibitory Smad7, and consequently suppressing TGF-β1/Smad signaling and HSC activation [42]. Similarly, Ki et al. demonstrated that SAHA improved collagen deposition by inhibiting HSC proliferation, inducing HSC apoptosis, and suppressing HSC activation in bile duct ligation (BDL)-induced cholestatic liver fibrosis model [74]. The recent study by Guan et al. demonstrated that SAHA suppresses TGF-β-induced chromatin structural changes and redirects mesenchymal cell differentiation away from activated myofibroblasts in human hepatic organoids [48]. In the current study, we complementarily focused on the protective effect of SAHA on LSECs, an aspect not addressed by previous studies. Together, prior studies and our findings suggest that the anti-fibrotic activity of SAHA may involve coordinated modulation of the LSEC-HSC fibrogenic unit: suppression of HSC activation on one side and restoration of endothelial-protective LSEC programs on the other. In addition to SAHA, multiple HDAC inhibitors have been shown to attenuate HSC activation and reduce collagen deposition across various experimental liver fibrosis models (e.g., HNHA in BDL model, MC1568 in CCl_4_ model, and valproate in TAA model) [75], strengthening the rationale for HDAC inhibitors selected by our platform (SAHA, mocetinostat, and resminostat) for treating liver fibrosis.

Beyond hepatotoxin-induced liver injury, the improvement of liver fibrosis in our established hepatocyte-specific *Asah1*-deficient PD model indicated that SAHA also protects LSECs under metabolic injury, especially metabolic stress associated with disrupted sphingolipid metabolism. Notably, SAHA did not improve hepatic steatosis and lipid concentrations, indicating that it may not directly target metabolic pathways. This suggests that therapeutic modulation of LSEC dysfunction could complement existing metabolic therapies (e.g., resmetirom, semaglutide) by targeting endothelial-associated fibrogenic processes that are not directly addressed by metabolic pathway modulation.

Another finding of the current study is that HMGB1, which was previously implicated in hepatocyte-to-HSC communication[52], directly mediates hepatocyte-to-LSEC paracrine injury. Our receptor interaction analysis revealed that HMGB1 signaling in LSECs is primarily mediated through TLR4 (pro-inflammatory), THBD (anti-inflammatory), and CXCR4 (remodeling), rather than the classic HMGB1-RAGE axis in HSCs[52]. In line with our findings, previous studies have reported the significant functions of HMGB1 with these three receptors in endothelial cells. Jia et al. reported that HMGB1 directly combined with TLR4 on LSECs, promoted IRF1 nuclear translocation, and finally led to the chemotaxis of neutrophils[76]. Nakamura et al. demonstrated that recombinant soluble THBD preserved LSECs in a rat model of sinusoidal obstruction syndrome by sequestering circulating HMGB1 and suppressing neutrophil recruitment[77]. Zhang et al. showed that HMGB1 mediates mitochondrial and endothelial function via CXCR4/PSMB5 pathway-mediated Drp1 proteolysis[78]. This LSEC-specific receptor landscape may contribute to future design of HMGB1 signaling- or LSEC-targeted interventions.

By analyzing the public dataset (GSE54912), we identified the endothelial transcription factor profile mediated by SAHA. Among these TFs, KLF2 upregulation by SAHA was confirmed in primary LSECs and in vivo. KLF2 activation is closely associated with anti-inflammation effects, including suppression on VCAM-1, CCL2, IL-6, and NLRP3 inflammasome components, all of which were suppressed by SAHA in primary LSECs, indicating the potential involvement of KLF2 signaling[79–81]. Another established vasoprotective effect of KLF2 is regulating vasodilation by promoting eNOS/NO signaling and suppressing ET-1 (encoded by EDN1) expression [79, 81]. In the current study, we did not observe significant increase of eNOS after SAHA treatment. Given that SAHA is a pan-HDAC inhibitor and modulates a broad TF profile, the effect on LSECs is likely mediated by the coordinated action of multiple TFs, including but not limited to *KLF2*. Therefore, unchanged eNOS expression may be a net effect of co-regulation by multiple TFs.

Nonetheless, SAHA significantly suppressed ET-1 expression. ET-1 has been reported as a potent vasoconstrictor that enhances the contractile response of HSCs, elevating sinusoidal vascular tone and exacerbating portal hypertension[82]. Additionally, ET-1 disrupts the balance between pro-fibrotic and anti-fibrotic factors, promoting excessive extracellular matrix synthesis, and thereby increases liver stiffness. Therefore, ET-1 suppression by SAHA may independently contribute to the improved portal hemodynamics observed in the *Asah1*-PD model. The lack of hemodynamic improvement in the CCl_4_ model may reflect differences in disease mechanisms (metabolic liver fibrosis and chemical-induced fibrosis) [83] and the distinct hemodynamic pathology (which may also account for inconsistent changes in PI).

KLF2 is also known to inhibit vascular endothelial growth factor (VEGF) signaling and *KDR* (encoding vascular endothelial growth factor receptor 2 [VEGFR2]) expression, and thereby suppress pathogenic angiogenesis [79, 81, 84]. However, VEGF signaling is also critical for maintaining the fenestrated structure of LSEC, and downregulation of VEGFR2 has been associated with LSEC capillarization in cirrhotic liver[9]. In the current study, SAHA suppressed basal expression of LYVE1 and KDR in primary LSECs in a dose-dependent manner, with significant reductions observed at medium (5 μM) and high (10 μM) concentrations, suggesting that SAHA may independently promote LSEC dedifferentiation and loss of fenestrae at higher doses[85]. However, this suppression was not observed in hepatic tissue, which may be attributed to the lower effective hepatic concentration achieved with systemic administration (15 mg/kg/day, i.p.) compared to direct in vitro exposure. This dose-dependent effect highlights the importance of therapeutic window optimization in future studies, and direct observation of LSEC fenestrae structure by electron microscopy. In addition to potential LSEC dedifferentiation at higher concentrations, clinical translation of systemic SAHA would require careful consideration of known dose-limiting toxicities, including thrombocytopenia, which is particularly relevant in patients with advanced liver fibrosis or cirrhosis [86, 87]. Therefore, future studies should define exposure-response relationships, LSEC target engagement, platelet effects, and whether endothelial-targeted delivery can preserve LSEC-protective activity while reducing systemic toxicity.

Another clinical-stage drug candidate identified by our platform, Amlexanox (TBK1/IKK-ε inhibitor), is particularly noteworthy because an independent Phase II clinical trial demonstrated improved hepatic steatosis in a subset of obese T2DM patients with concurrent NAFLD[40]. TBK1/IKKε are both pro-inflammatory and pro-fibrotic kinases upregulated in fibrotic livers and activated HSCs, indicating the potential anti-inflammatory and anti-fibrotic effects of Amlexanox [88]. However, this trial was not designed for NAFLD treatment and thus did not include hepatic inflammation or fibrosis as endpoints, which may underestimate its repurposing potential. Additionally, a recent study reported that Amlexanox reversed MASH and prevented HCC progression in GAN diet-fed *Ldlr*^-/-^ mice by promoting bile acid synthesis and fecal bile acid excretion, suggesting regulation of bile acid metabolism[41]. Therefore, Amlexanox’s repurposing potential in fibrotic MASH and cholestatic liver diseases may be worth future study.

Together, the in vivo, co-culture, and primary LSEC validation studies support a proposed working model in which SAHA preferentially attenuates hepatic inflammation, fibrosis, LSEC injury, and portal venous velocity impairment in experimental liver fibrosis, while hepatic steatosis, cardiac dysfunction, and potential dose-dependent effects on LSEC differentiation markers remain unresolved **(Figure 10).**

**Figure 10.**
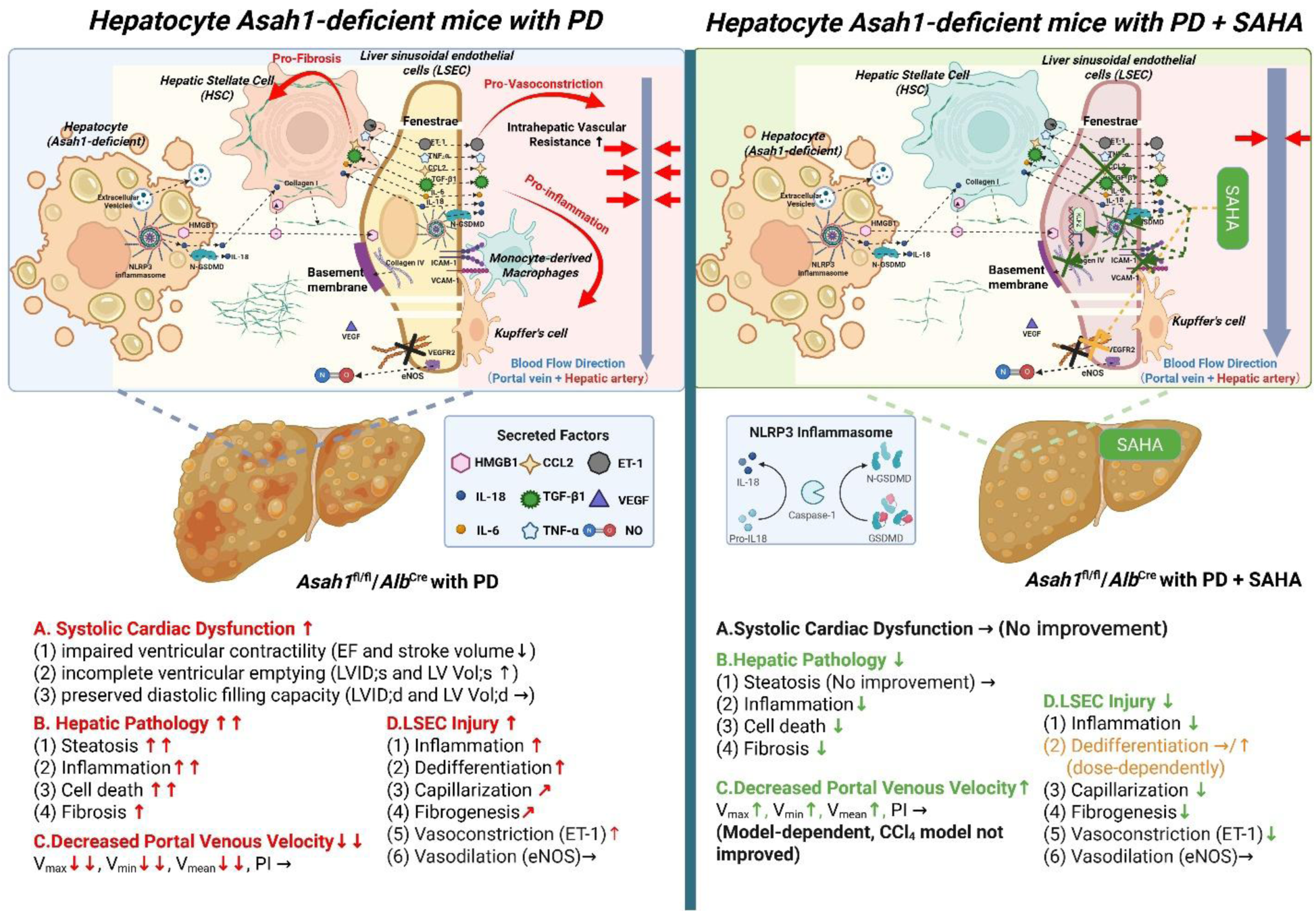
Proposed working model of SAHA-mediated LSEC protection in experimental liver fibrosis. This schematic summarizes the major findings of this study and presents a proposed working model. In hepatocyte-specific *Asah1*-deficient mice fed a Paigen diet, hepatocyte injury and paracrine signaling, including HMGB1 release, are associated with hepatic inflammation, fibrosis, LSEC dysfunction, reduced portal venous velocity, and systolic cardiac dysfunction. SAHA attenuates hepatic inflammation, fibrosis, LSEC inflammatory activation, capillarization-associated changes, fibrogenic signaling, and ET-1-associated vasoconstrictive signaling, and improves portal venous velocity. Similar anti-fibrotic and LSEC-protective effects were observed in the CCl4-induced fibrosis model, although portal venous hemodynamics were not significantly improved. SAHA does not improve hepatic steatosis or systolic cardiac dysfunction, and in vitro findings suggest potential dose-dependent effects on LSEC differentiation markers, highlighting the need for therapeutic window optimization. Abbreviations: PD, Paigen diet; HSC, hepatic stellate cell; LSEC, liver sinusoidal endothelial cell; KC, Kupffer cell; CCl_4_, carbon tetrachloride; CCL2, C-C motif chemokine ligand 2; ET-1, endothelin-1; GSDMD, gasdermin D; HMGB1, high mobility group box 1; IL-6, interleukin-6; IL-18, interleukin-18; N-GSDMD, N-terminal gasdermin D; NLRP3, NOD-, LRR- and pyrin domain-containing protein 3; NO, nitric oxide; Pro-IL18, pro-interleukin-18; TGF-β1, transforming growth factor beta 1; TNF-α, tumor necrosis factor alpha; VEGF, vascular endothelial growth factor; ICAM-1, intercellular adhesion molecule 1; VCAM-1, vascular cell adhesion molecule 1; eNOS, endothelial nitric oxide synthase; EF, ejection fraction; LV Vol;d, left ventricular volume in diastole; LV Vol;s, left ventricular volume in systole; LVID;d, left ventricular internal diameter in diastole; LVID;s, left ventricular internal diameter in systole; PI, pulsatility index; SAHA, suberoylanilide hydroxamic acid/vorinostat; Vmax, maximum velocity; Vmean, mean velocity; Vmin, minimum velocity.

Several limitations should be acknowledged. First, the CDAA diet used in GSE140994 does not recapitulate the metabolic features of human MASLD/MASH (e.g., insulin resistance, obesity)[89]. Second, SAHA did not improve cardiac function in the hepatocyte-specific *Asah1*-deficient PD model, which may be partially attributed to the first-pass effect, or the involvement of liver-derived mediators that are not targeted by SAHA. For example, PA-treated hepatocytes have been demonstrated to release C16:0 ceramide-enriched extracellular vesicles (EVs), activating macrophage chemotaxis[65], where circulating ceramide-enriched EV is closely related to cardiovascular events[90]. Considering that *ASAH1* is responsible for ceramide degradation, ceramide-mediated liver-to-heart crosstalk may contribute to cardiac dysfunction independently of hepatic fibrosis. In addition, the mechanism study of SAHA on LSEC protection remains preliminary. Future studies will involve systematic characterization of SAHA’s epigenomic effects in LSECs, including HDAC isoform-selective profiling and chromatin accessibility analysis (e.g., assay for transposase-accessible chromatin with sequencing, [ATAC-seq]). LSEC-specific genetic models (e.g., *Cdh5*-Cre or *Stab2*-Cre driven systems) targeting key endothelial-protective TFs (such as KLF2) will further clarify their roles. Another limitation is that the direct endothelial-protective effects of SAHA were validated primarily in mouse primary LSECs. Although human endothelial perturbational data and human cirrhotic LSEC signatures were incorporated computationally, future studies using primary human LSECs or human LSEC-like models will be needed to further establish human relevance. Moreover, although co-immunofluorescence and primary LSEC experiments support an LSEC-protective effect, the current in vivo studies do not prove that LSECs are the required causal cellular target of SAHA. LSEC-specific genetic or pharmacologic perturbation studies will be needed to define the contribution of endothelial mechanisms relative to effects on HSCs, immune cells, and hepatocytes.

In conclusion, this study establishes a computational drug screening framework focused on LSECs, an underestimated cell type in anti-fibrotic drug development. The platform identified a series of clinical and preclinical drug candidates with LSEC-protective potential across multiple classes, with independent clinical/preclinical evidence supporting translational potential. As proof-of-concept, SAHA demonstrated anti-fibrotic and LSEC-protective effects in two independent mouse models, associated with endothelial TF reprogramming. These findings support LSEC dysfunction as a tractable therapeutic axis in liver fibrosis and provide a framework for future cell-type-focused drug repurposing and translational development.

## DATA AVAILABILITY STATEMENT

Publicly available scRNA-seq and bulk RNA-seq datasets were obtained from the GEO dataset (NCBI, https://www.ncbi.nlm.nih.gov/gds) under accession numbers GSE136103, GSE140994, GSE166504, and GSE54912. All other data and code are available from the corresponding author upon reasonable request.

## AI USAGE STATEMENT

Claude Opus 4.0 (Anthropic) was used for gene signature curation of the computational platform, as described in **Methods**. Additionally, AI-assisted tools were used for language and code polishing. All AI-involved content was reviewed and approved by the authors, who take full responsibility for the work.

## ACKNOWLEDGEMENTS

This study was supported by NIH grants R01HL122937 (Y.Z.) and R01HL150007 (X.L.).

## SUPPLEMENTARY MATERIALS

**Supplementary Figure 1.**
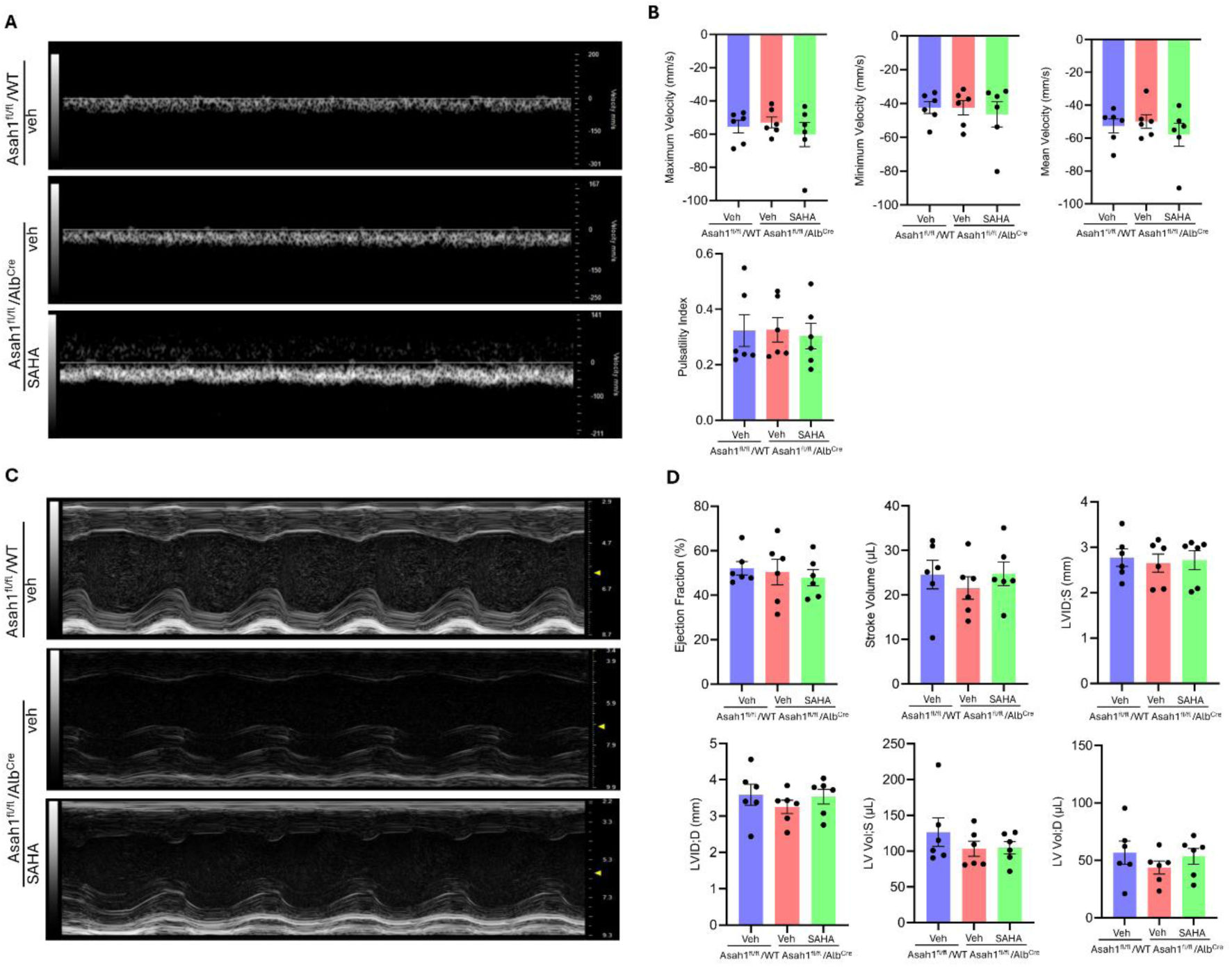
Portal hemodynamics and cardiac function are preserved at week 12 in hepatocyte-specific *Asah1* deficient mice. *Asah1*^fl/fl^/*Alb*^Cre^ mice (hepatocyte-specific deletion of *Asah1*) and *Asah1* floxed (*Asah1*^fl/fl^/WT) mice were fed with PD from week 0, treated with vehicle or SAHA (15 mg/kg/day, i.p.) from week 6. Portal venous hemodynamics and cardiac function were assessed by ultrasonography at week 12. **(A-B)** Representative portal vein Doppler waveforms and hemodynamic quantification including maximum velocity (mm/s), minimum velocity (mm/s), mean velocity (mm/s), and pulsatility index. **(C-D)** Representative echocardiography images and cardiac function parameters including Ejection Fraction (%), Stroke Volume (μL), LVID D (mm), LVID S (mm), LV Vol D (μL), and LV Vol S (μL). n = 5-6/group. Abbreviations: LVID D, left ventricular internal diameter in diastole; LVID S, left ventricular internal diameter in systole; LV Vol D, left ventricular volume in diastole; LV Vol S, left ventricular volume in systole.

**Supplementary Table 1.**
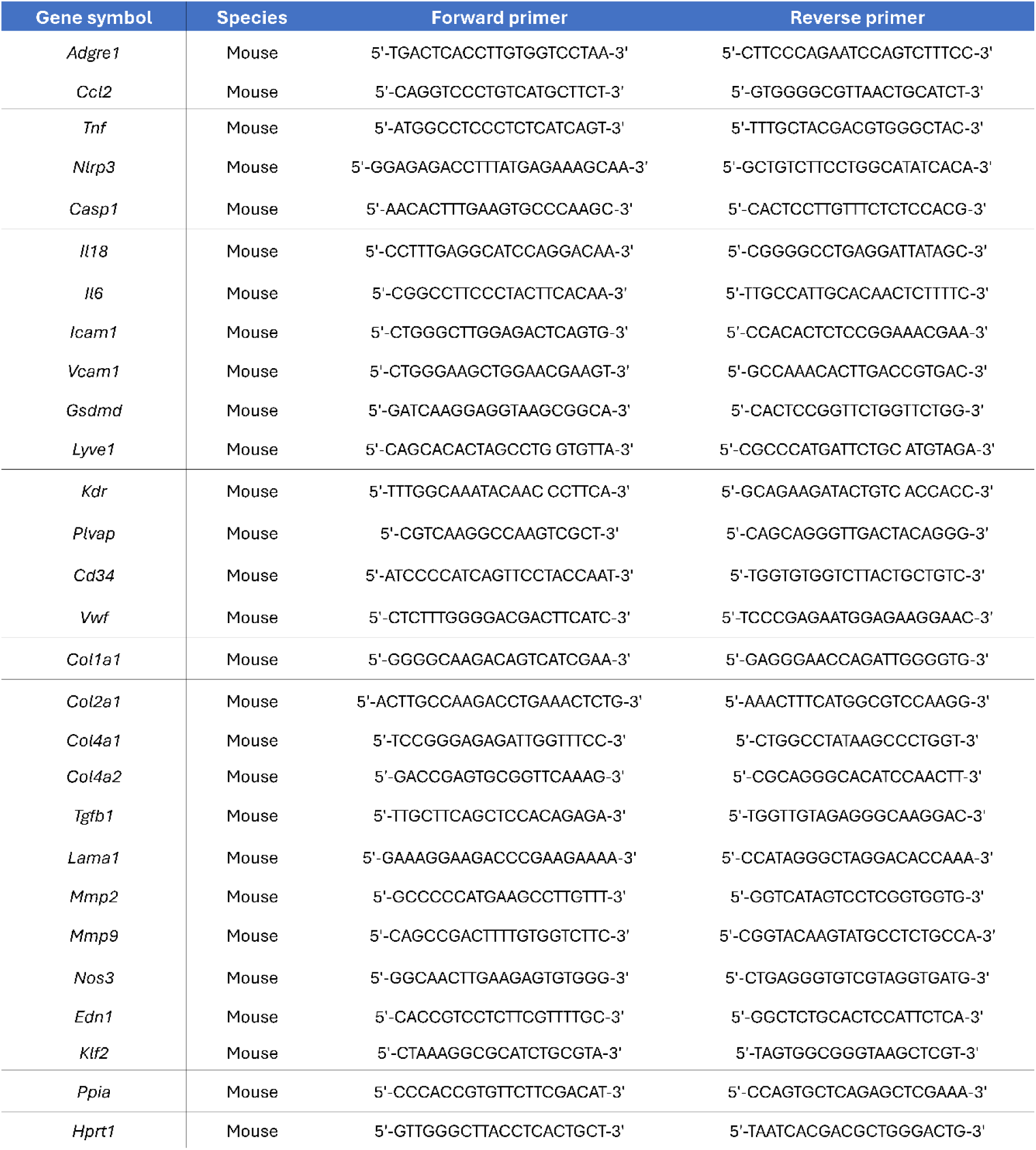
Primer Sequences.

**Supplementary Table 2.**
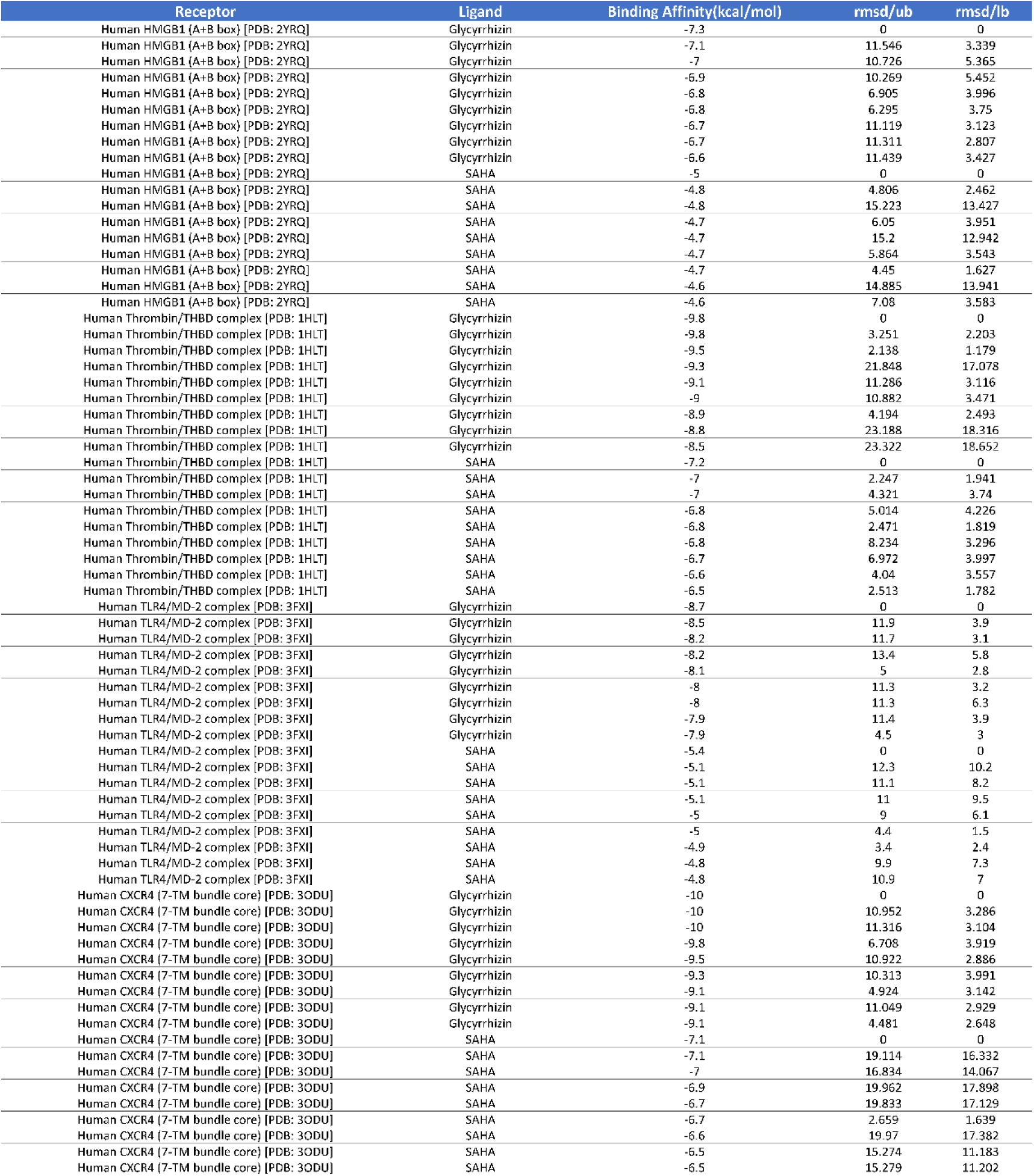
Molecular docking results of Gly and SAHA against HMGB1 and its LSEC receptors. Binding affinities (kcal/mol) and RMSD values for Gly and SAHA docked against four protein targets: HMGB1 A+B box (PDB: 2YRQ), thrombin/THBD complex (PDB: 1HLT), TLR4/MD-2 complex (PDB: 3FXI), and CXCR4 7-TM bundle core (PDB: 3ODU). Up to ten docking poses per ligand-receptor pair were generated using AutoDock Vina. More negative binding affinity values indicate stronger predicted binding. RMSD/ub, upper bound root-mean-square deviation; RMSD/lb, lower bound root-mean-square deviation. The top-ranked pose (RMSD = 0) for each ligand-receptor pair was used for interaction analysis in Figure 7. Abbreviations: Gly, glycyrrhizin; PDB, Protein Data Bank; THBD, thrombomodulin; TLR4, Toll-like receptor 4; MD-2, myeloid differentiation factor 2; CXCR4, C-X-C chemokine receptor type 4; 7-TM, seven-transmembrane; RMSD, root-mean-square deviation.

**Supplementary Table 3.**
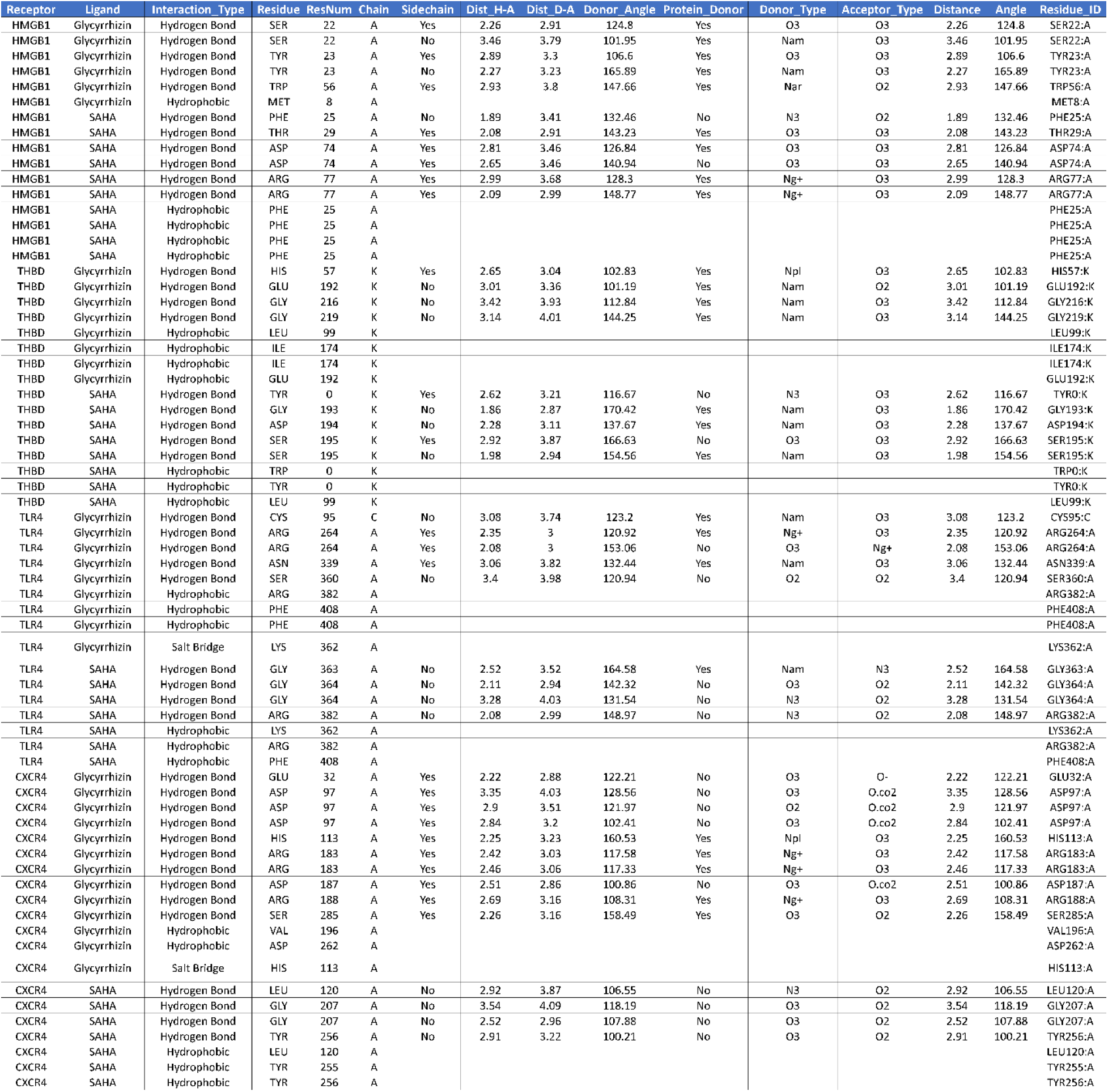
Detailed molecular interaction profiles of Gly and SAHA with HMGB1 and its LSEC receptors. Residue-level interaction analysis for Gly and SAHA docked against HMGB1 (PDB: 2YRQ), THBD (PDB: 1HLT), TLR4/MD-2 (PDB: 3FXI), and CXCR4 (PDB: 3ODU). Interaction profiles were generated using PLIP (Protein-Ligand Interaction Profiler) from top-ranked docking poses **(Supplementary Table 2)**. Interaction types include hydrogen bonds, hydrophobic interactions, and salt bridges. For hydrogen bonds, donor-acceptor distances (Dist H-A, Dist D-A), angles (Donor_Angle, Protein_Donor), donor/acceptor types, and sidechain involvement are reported. Abbreviations: Gly, glycyrrhizin; THBD, thrombomodulin; TLR4, Toll-like receptor 4; MD-2, myeloid differentiation factor 2; CXCR4, C-X-C chemokine receptor type 4; PDB, Protein Data Bank; PLIP, Protein-Ligand Interaction Profiler; ResNum, residue number; Dist H-A, hydrogen-acceptor distance; Dist D-A, donor-acceptor distance.

## REFERENCES

1. Roehlen, N., E. Crouchet, and T.F. Baumert, Liver Fibrosis: Mechanistic Concepts and Therapeutic Perspectives. Cells, 2020. 9(4).

2. Devarbhavi, H., et al., Global burden of liver disease: 2023 update. J Hepatol, 2023. 79(2): p. 516–537.

3. Zamani, M., et al., Global Prevalence of Advanced Liver Fibrosis and Cirrhosis in the General Population: A Systematic Review and Meta-analysis. Clin Gastroenterol Hepatol, 2025. 23(7): p. 1123–1134.

4. Harrison, S.A., et al., Resmetirom (MGL-3196) for the treatment of non-alcoholic steatohepatitis: a multicentre, randomised, double-blind, placebo-controlled, phase 2 trial. Lancet, 2019. 394(10213): p. 2012–2024.

5. Loomba, R., et al., Semaglutide 2·4 mg once weekly in patients with non-alcoholic steatohepatitis-related cirrhosis: a randomised, placebo-controlled phase 2 trial. Lancet Gastroenterol Hepatol, 2023. 8(6): p. 511–522.

6. Tsuchida, T. and S.L. Friedman, Mechanisms of hepatic stellate cell activation. Nat Rev Gastroenterol Hepatol, 2017. 14(7): p. 397–411.

7. Mao, X., et al., From mechanisms to anti-fibrotic drugs in hepatic stellate cell research: a global bibliometric analysis with patent and clinical perspectives (2000-2025). Front Pharmacol, 2025. 16: p. 1734449.

8. Qu, J., et al., Liver sinusoidal endothelial cell: An important yet often overlooked player in the liver fibrosis. Clin Mol Hepatol, 2024. 30(3): p. 303–325.

9. Iwakiri, Y., Unlocking the role of liver sinusoidal endothelial cells: Key players in liver fibrosis: Editorial on “Liver sinusoidal endothelial cell: An important yet often overlooked player in the liver fibrosis”. Clin Mol Hepatol, 2024. 30(4): p. 673–676.

10. Felli, E., et al., The role of liver sinusoidal endothelial cells in liver diseases: Key players in health and pathology. Journal of Hepatology, 2026. 84(3): p. 655–672.

11. Gracia-Sancho, J., et al., Role of liver sinusoidal endothelial cells in liver diseases. Nat Rev Gastroenterol Hepatol, 2021. 18(6): p. 411–431.

12. Marrone, G., et al., The transcription factor KLF2 mediates hepatic endothelial protection and paracrine endothelial-stellate cell deactivation induced by statins. J Hepatol, 2013. 58(1): p. 98–103.

13. DeLeve, L.D., Liver sinusoidal endothelial cells in hepatic fibrosis. Hepatology, 2015. 61(5): p. 1740–6.

14. Miyao, M., et al., Pivotal role of liver sinusoidal endothelial cells in NAFLD/NASH progression. Lab Invest, 2015. 95(10): p. 1130–44.

15. Xie, G., et al., Role of differentiation of liver sinusoidal endothelial cells in progression and regression of hepatic fibrosis in rats. Gastroenterology, 2012. 142(4): p. 918–927.e6.

16. Hu, Y., et al., Selective targeting of endothelial and perivascular angiocrine ROCK2 treats liver fibrosis. Cell, 2026. 189(9): p. 2663–2683.e26.

17. Lamb, J., et al., The Connectivity Map: using gene-expression signatures to connect small molecules, genes, and disease. Science, 2006. 313(5795): p. 1929–35.

18. Su, Q., et al., Single-cell RNA transcriptome landscape of hepatocytes and non-parenchymal cells in healthy and NAFLD mouse liver. iScience, 2021. 24(11): p. 103233.

19. Winkler, M., et al., Endothelial GATA4 controls liver fibrosis and regeneration by preventing a pathogenic switch in angiocrine signaling. J Hepatol, 2021. 74(2): p. 380–393.

20. Ramachandran, P., et al., Resolving the fibrotic niche of human liver cirrhosis at single-cell level. Nature, 2019. 575(7783): p. 512–518.

21. Hao, Y., et al., Dictionary learning for integrative, multimodal and scalable single-cell analysis. Nat Biotechnol, 2024. 42(2): p. 293–304.

22. Robinson, M.D., D.J. McCarthy, and G.K. Smyth, edgeR: a Bioconductor package for differential expression analysis of digital gene expression data. Bioinformatics, 2010. 26(1): p. 139–40.

23. Ritchie, M.E., et al., limma powers differential expression analyses for RNA-sequencing and microarray studies. Nucleic Acids Res, 2015. 43(7): p. e47.

24. Hänzelmann, S., R. Castelo, and J. Guinney, GSVA: gene set variation analysis for microarray and RNA-seq data. BMC Bioinformatics, 2013. 14: p. 7.

25. Liberzon, A., et al., The Molecular Signatures Database (MSigDB) hallmark gene set collection. Cell Syst, 2015. 1(6): p. 417–425.

26. Yu, G., et al., clusterProfiler: an R package for comparing biological themes among gene clusters. Omics, 2012. 16(5): p. 284–7.

27. Meinshausen, N. and P. Bühlmann, Stability Selection. Journal of the Royal Statistical Society Series B: Statistical Methodology, 2010. 72(4): p. 417–473.

28. Friedman, J., T. Hastie, and R. Tibshirani, Regularization Paths for Generalized Linear Models via Coordinate Descent. J Stat Softw, 2010. 33(1): p. 1–22.

29. Tsherniak, A., et al., Defining a Cancer Dependency Map. Cell, 2017. 170(3): p. 564–576.e16.

30. Sondka, Z., et al., The COSMIC Cancer Gene Census: describing genetic dysfunction across all human cancers. Nat Rev Cancer, 2018. 18(11): p. 696–705.

31. Landrum, M.J., et al., ClinVar: improving access to variant interpretations and supporting evidence. Nucleic Acids Res, 2018. 46(D1): p. D1062–d1067.

32. Subramanian, A., et al., A Next Generation Connectivity Map: L1000 Platform and the First 1,000,000 Profiles. Cell, 2017. 171(6): p. 1437–1452.e17.

33. Dallakyan, S. and A.J. Olson, Small-molecule library screening by docking with PyRx. Methods Mol Biol, 2015. 1263: p. 243–50.

34. Trott, O. and A.J. Olson, AutoDock Vina: improving the speed and accuracy of docking with a new scoring function, efficient optimization, and multithreading. J Comput Chem, 2010. 31(2): p. 455–61.

35. Adasme, M.F., et al., PLIP 2021: expanding the scope of the protein-ligand interaction profiler to DNA and RNA. Nucleic Acids Res, 2021. 49(W1): p. W530–w534.

36. Zuo, R., et al., Ablation of Hepatic Asah1 Gene Disrupts Hepatic Lipid Homeostasis and Promotes Fibrotic Nonalcoholic Steatohepatitis in Mice. Am J Pathol, 2025. 195(3): p. 542–560.

37. Wang, Y.T., et al., Cardiovascular dysfunction and altered lysosomal signaling in a murine model of acid sphingomyelinase deficiency. J Mol Med (Berl), 2025. 103(5): p. 599–617.

38. Wang, Y.T., et al., Coronary Microvascular Dysfunction Is Associated With Augmented Lysosomal Signaling in Hypercholesterolemic Mice. J Am Heart Assoc, 2024. 13(23): p. e037460.

39. Szklarczyk, D., et al., The STRING database in 2023: protein-protein association networks and functional enrichment analyses for any sequenced genome of interest. Nucleic Acids Res, 2023. 51(D1): p. D638–d646.

40. Oral, E.A., et al., Inhibition of IKKɛ and TBK1 Improves Glucose Control in a Subset of Patients with Type 2 Diabetes. Cell Metab, 2017. 26(1): p. 157–170.e7.

41. You, W., et al., Unlocking therapeutic potential of amlexanox in MASH with insights into bile acid metabolism and microbiome. NPJ Gut Liver, 2025. 2.

42. Wang, Y., et al., Histone deacetylase inhibitor suberoylanilide hydroxamic acid alleviates liver fibrosis by suppressing the transforming growth factor-β1 signal pathway. Hepatobiliary Pancreat Dis Int, 2018. 17(5): p. 423–429.

43. Bitzer, M., et al., Resminostat plus sorafenib as second-line therapy of advanced hepatocellular carcinoma - The SHELTER study. J Hepatol, 2016. 65(2): p. 280–8.

44. Middleton, S.A., et al., BET Inhibition Improves NASH and Liver Fibrosis. Sci Rep, 2018. 8(1): p. 17257.

45. Foglia, B., et al., ERK Pathway in Activated, Myofibroblast-Like, Hepatic Stellate Cells: A Critical Signaling Crossroad Sustaining Liver Fibrosis. Int J Mol Sci, 2019. 20(11).

46. Zhong, W., et al., Inhibition of extracellular signal-regulated kinase 1 by adenovirus mediated small interfering RNA attenuates hepatic fibrosis in rats. Hepatology, 2009. 50(5): p. 1524–36.

47. Zhang, Y.B., et al., Hydroxysafflor yellow A attenuates carbon tetrachloride-induced hepatic fibrosis in rats by inhibiting ERK5 signaling. Am J Chin Med, 2012. 40(3): p. 481–94.

48. Guan, Y., et al., AI-Assisted Drug Re-Purposing for Human Liver Fibrosis. Adv Sci (Weinh), 2025. 12(44): p. e08751.

49. Beraza, N., et al., Pharmacological IKK2 inhibition blocks liver steatosis and initiation of non-alcoholic steatohepatitis. Gut, 2008. 57(5): p. 655–63.

50. Liang, D., et al., Inhibition of EGFR attenuates fibrosis and stellate cell activation in diet-induced model of nonalcoholic fatty liver disease. Biochim Biophys Acta Mol Basis Dis, 2018. 1864(1): p. 133–142.

51. Wang, F., et al., Canonical Wnt signaling promotes HSC glycolysis and liver fibrosis through an LDH-A/HIF-1α transcriptional complex. Hepatology, 2024. 79(3): p. 606–623.

52. Ge, X., et al., High Mobility Group Box-1 Drives Fibrosis Progression Signaling via the Receptor for Advanced Glycation End Products in Mice. Hepatology, 2018. 68(6): p. 2380–2404.

53. Kakisaka, K., et al., A hedgehog survival pathway in ‘undead’ lipotoxic hepatocytes. J Hepatol, 2012. 57(4): p. 844–51.

54. Rangwala, F., et al., Increased production of sonic hedgehog by ballooned hepatocytes. J Pathol, 2011. 224(3): p. 401–10.

55. Chung, S.I., et al., Hepatic expression of Sonic Hedgehog induces liver fibrosis and promotes hepatocarcinogenesis in a transgenic mouse model. J Hepatol, 2016. 64(3): p. 618–27.

56. Wang, X., et al., Hepatocyte TAZ/WWTR1 Promotes Inflammation and Fibrosis in Nonalcoholic Steatohepatitis. Cell Metab, 2016. 24(6): p. 848–862.

57. Yan, T., et al., Hepatocyte-specific CCAAT/enhancer binding protein α restricts liver fibrosis progression. J Clin Invest, 2024. 134(7).

58. Urtasun, R., et al., Osteopontin, an oxidant stress sensitive cytokine, up-regulates collagen-I via integrin α(V)β(3) engagement and PI3K/pAkt/NFκB signaling. Hepatology, 2012. 55(2): p. 594–608.

59. Dong, J., et al., Hepatocyte-specific IL11 cis-signaling drives lipotoxicity and underlies the transition from NAFLD to NASH. Nat Commun, 2021. 12(1): p. 66.

60. Povero, D., et al., Lipid-induced hepatocyte-derived extracellular vesicles regulate hepatic stellate cell via microRNAs targeting PPAR-γ. Cell Mol Gastroenterol Hepatol, 2015. 1(6): p. 646–663.e4.

61. Yan, Z., et al., CD147 promotes liver fibrosis progression via VEGF-A/VEGFR2 signalling-mediated cross-talk between hepatocytes and sinusoidal endothelial cells. Clin Sci (Lond), 2015. 129(8): p. 699–710.

62. Abad-Jordà, L., et al., Hepatocyte-derived extracellular vesicles promote endothelial dedifferentiation in chronic liver disease through the miR-153-3p-pyroptosis axis. Hepatology, 2025.

63. Guo, Q., et al., Integrin β(1)-enriched extracellular vesicles mediate monocyte adhesion and promote liver inflammation in murine NASH. J Hepatol, 2019. 71(6): p. 1193–1205.

64. Kang, J., et al., Notch-mediated hepatocyte MCP-1 secretion causes liver fibrosis. JCI Insight, 2023. 8(3).

65. Kakazu, E., et al., Hepatocytes release ceramide-enriched pro-inflammatory extracellular vesicles in an IRE1α-dependent manner. J Lipid Res, 2016. 57(2): p. 233–45.

66. Liu, X.L., et al., Lipotoxic Hepatocyte-Derived Exosomal MicroRNA 192-5p Activates Macrophages Through Rictor/Akt/Forkhead Box Transcription Factor O1 Signaling in Nonalcoholic Fatty Liver Disease. Hepatology, 2020. 72(2): p. 454–469.

67. Ibrahim, S.H., et al., Mixed lineage kinase 3 mediates release of C-X-C motif ligand 10-bearing chemotactic extracellular vesicles from lipotoxic hepatocytes. Hepatology, 2016. 63(3): p. 731–44.

68. Tomita, K., et al., CXCL10-Mediates Macrophage, but not Other Innate Immune Cells-Associated Inflammation in Murine Nonalcoholic Steatohepatitis. Sci Rep, 2016. 6: p. 28786.

69. Hirsova, P., et al., Lipid-Induced Signaling Causes Release of Inflammatory Extracellular Vesicles From Hepatocytes. Gastroenterology, 2016. 150(4): p. 956–67.

70. Garcia-Martinez, I., et al., Hepatocyte mitochondrial DNA drives nonalcoholic steatohepatitis by activation of TLR9. J Clin Invest, 2016. 126(3): p. 859–64.

71. Valenti, L., S. Romeo, and U. Pajvani, A genetic hypothesis for burnt-out steatohepatitis. Liver Int, 2021. 41(12): p. 2816–2818.

72. Hédou, J., et al., Discovery of sparse, reliable omic biomarkers with Stabl. Nat Biotechnol, 2024. 42(10): p. 1581–1593.

73. Moon, M. and K. Nakai, Stable feature selection based on the ensemble L (1) -norm support vector machine for biomarker discovery. BMC Genomics, 2016. 17(Suppl 13): p. 1026.

74. Park, K.C., et al., A new histone deacetylase inhibitor improves liver fibrosis in BDL rats through suppression of hepatic stellate cells. Br J Pharmacol, 2014. 171(21): p. 4820–30.

75. Yoon, S., G. Kang, and G.H. Eom, HDAC Inhibitors: Therapeutic Potential in Fibrosis-Associated Human Diseases. Int J Mol Sci, 2019. 20(6).

76. Jia, K., et al., Acteoside ameliorates hepatic ischemia-reperfusion injury via reversing the senescent fate of liver sinusoidal endothelial cells and restoring compromised sinusoidal networks. Int J Biol Sci, 2023. 19(15): p. 4967–4988.

77. Nakamura, K., et al., Soluble thrombomodulin attenuates sinusoidal obstruction syndrome in rat through suppression of high mobility group box 1. Liver Int, 2014. 34(10): p. 1473–87.

78. Zhang, S., et al., Recombinant High-Mobility Group Box 1 (rHMGB1) Promotes NRF2-Independent Mitochondrial Fusion through CXCR4/PSMB5-Mediated Drp1 Degradation in Endothelial Cells. Oxid Med Cell Longev, 2021. 2021: p. 9993240.

79. Nayak, L., Z. Lin, and M.K. Jain, ”Go with the flow”: how Krüppel-like factor 2 regulates the vasoprotective effects of shear stress. Antioxid Redox Signal, 2011. 15(5): p. 1449–61.

80. Zhuang, T., et al., Endothelial Foxp1 Suppresses Atherosclerosis via Modulation of Nlrp3 Inflammasome Activation. Circ Res, 2019. 125(6): p. 590–605.

81. Dekker, R.J., et al., KLF2 provokes a gene expression pattern that establishes functional quiescent differentiation of the endothelium. Blood, 2006. 107(11): p. 4354–63.

82. Ezhilarasan, D., Endothelin-1 in portal hypertension: The intricate role of hepatic stellate cells. Exp Biol Med (Maywood), 2020. 245(16): p. 1504–1512.

83. Wu, S., et al., An update on animal models of liver fibrosis. Front Med (Lausanne), 2023. 10: p. 1160053.

84. Bhattacharya, R., et al., Inhibition of vascular permeability factor/vascular endothelial growth factor-mediated angiogenesis by the Kruppel-like factor KLF2. J Biol Chem, 2005. 280(32): p. 28848–51.

85. Su, T., et al., Single-Cell Transcriptomics Reveals Zone-Specific Alterations of Liver Sinusoidal Endothelial Cells in Cirrhosis. Cell Mol Gastroenterol Hepatol, 2021. 11(4): p. 1139–1161.

86. Afdhal, N., et al., Thrombocytopenia associated with chronic liver disease. J Hepatol, 2008. 48(6): p. 1000–7.

87. Ramalingam, S.S., et al., Phase I study of vorinostat in patients with advanced solid tumors and hepatic dysfunction: a National Cancer Institute Organ Dysfunction Working Group study. J Clin Oncol, 2010. 28(29): p. 4507–12.

88. Zhou, Z., et al., Dual TBK1/IKKɛ inhibitor amlexanox attenuates the severity of hepatotoxin-induced liver fibrosis and biliary fibrosis in mice. J Cell Mol Med, 2020. 24(2): p. 1383–1398.

89. Wei, G., et al., Comparison of murine steatohepatitis models identifies a dietary intervention with robust fibrosis, ductular reaction, and rapid progression to cirrhosis and cancer. Am J Physiol Gastrointest Liver Physiol, 2020. 318(1): p. G174–g188.

90. Timmerman, N., et al., Ceramides and phospholipids in plasma extracellular vesicles are associated with high risk of major cardiovascular events after carotid endarterectomy. Sci Rep, 2022. 12(1): p. 5521.

